# Identification of Antimalarial Compounds that Inhibit Apicomplexan AP2 Proteins in the Human Malaria Parasite *Plasmodium falciparum*

**DOI:** 10.1101/2022.04.05.487101

**Authors:** Timothy Russell, Erandi K. De Silva, Valerie Crowley, Kathryn Shaw-Saliba, Namita Dube, Gabrielle Josling, Charisse Flerida A. Pasaje, Irene Kouskoumvekaki, Gianni Panagiotou, Jacquin C. Niles, Marcelo Jacobs-Lorena, C. Denise Okafor, Francisco-Javier Gamo, Manuel Llinás

## Abstract

*Plasmodium* parasites are reliant on the Apicomplexan AP2 (ApiAP2) transcription factor family to regulate gene expression programs. AP2 DNA binding domains have no homologs in the human or mosquito host genomes, making them potential antimalarial drug targets. Using an *in-silico* screen to dock thousands of small molecules into the crystal structure of the AP2-EXP (Pf3D7_1466400) AP2 domain (PDB:3IGM), we identified compounds that interact with this domain. Four compounds were found to compete for DNA binding with AP2-EXP and at least one additional ApiAP2 protein. Our top ApiAP2 competitor compound perturbs the transcriptome of *P. falciparum* trophozoites and results in a decrease in abundance of log_2_ fold change > 2 for 50% (46/93) of AP2-EXP target genes. Additionally, two ApiAP2 competitor compounds have multi-stage anti-*Plasmodium* activity against blood and mosquito stage parasites. In summary, we describe a novel set of antimalarial compounds that are targeted against the ApiAP2 family of proteins. These compounds may be used for future chemical genetic interrogation of ApiAP2 proteins or serve as starting points for a new class of antimalarial therapeutics.

**Author Summary:** *Plasmodium* parasites are the causative agent of malaria, which resulted in over 600,000 deaths in 2021. Due to resistance arising for every antimalarial therapeutic deployed to date, new drug targets and druggable pathways must be explored. To address this concern, we used a molecular docking screen to predict competitors of DNA binding by the parasite specific family of Apicomplexan AP2 (ApiAP2) transcription factor proteins for testing *in vitro* and *in vivo*. We find that ApiAP2 competing compounds have antimalarial activity consistent with the disruption of gene regulation. This work will further our understanding of both the biological role and targetability of parasite transcriptional regulation.

## Introduction

Malaria is a disease caused by intracellular parasites from the genus *Plasmodium* that represents a significant health and economic burden worldwide(1). The most virulent of the human infectious malaria parasites is *Plasmodium falciparum*, which caused more than 200 million cases of malaria and resulted in over 600 thousand deaths in 2021(1). Resistance has been reported for every antimalarial therapeutic deployed to date, necessitating the need for malaria drugs that target new parasite processes(2). To successfully proliferate, *P. falciparum* parasites must develop through a complex lifecycle that includes intracellular and extracellular stages in both the human and *Anopheles* mosquito hosts(3). The clinical symptoms of malaria are caused by the intraerythrocytic development cycle (IDC), a 48-hour cyclic asexual proliferation that results in the destruction of red blood cells (RBCs). During the asexual IDC, malaria parasites progress through three morphological stages referred to as ring, trophozoite, and schizont. Parasites transmit from human to mosquito following differentiation and maturation into sexual stage gametocytes. Once ingested by the mosquito, gametes sexually reproduce and ultimately develop into sporozoites, which can be transmitted back to the human host to initiate a liver stage infection which precedes the asexual blood stages(3).

Up to 80% of *P. falciparum* protein coding transcripts are developmentally regulated during the IDC(4–8) as part of a ‘just in time’ cascade(5, 8, 9). This is predicted to be principally driven by the 27 Apicomplexan Apetala AP2 (ApiAP2) proteins(10–12), which are the major family of sequence specific transcription factors encoded in the *Plasmodium* genome. ApiAP2 proteins contain one to three AP2 DNA binding domains that are analogous to the plant APETALA2/Ethylene Responsive Factor (AP2/ERF) domain and therefore have no homologs in the human or mosquito genome(13, 14). Over half of the ApiAP2 proteins are predicted to be essential for the IDC in *P. falciparum*(15–17). In-depth studies of ApiAP2 proteins to date have demonstrated roles in transcriptional activation or repression affecting diverse processes including invasion(18, 19), heterochromatin maintenance(17, 20–23), sexual and mosquito stage differentiation(16, 24–34), and heat stress tolerance(35). Despite these properties which make ApiAP2 proteins potential drug targets, no efforts to date have focused on targeting the sequence specific DNA binding ApiAP2 transcription factors.

In this study we conducted an *in silico* chemical screen against the crystal structure of the AP2-EXP AP2 domain(36) to predict competitors of DNA binding activity. AP2-EXP is predicted to be essential for the *P. falciparum* asexual blood stage,(16, 17, 21) and its orthologue in the rodent malaria parasite *P. berghei* (PbAP2-Sp) is a master regulator of sporogony(32). Several compounds identified in the screen can compete for DNA binding activity with the AP2-EXP AP2 domain *in vitro*. We also find that modifications to the substitution state of an AP2 domain competitor compound can modulate its ability to compete DNA binding. Our lead compound kills *P. falciparum* parasites in concordance with the stage specific expression of AP2-EXP. We measured AP2-EXP genomic occupancy by Chromatin immunoprecipitation followed by high throughput sequencing (ChIP-seq) and found high correlation between dysregulated mRNA transcripts in the presence of our lead compound and AP2-EXP gene targets, corroborating the hypothesis that AP2-EXP is inhibited *in vivo*. Finally, two AP2 domain competitor compounds are active against *P. berghei* development in the mosquito stage. Overall, our results demonstrate the potential to chemically compete the DNA binding activity of ApiAP2 proteins.

## Results

### *In silico* Prediction of AP2-EXP Competitors

To identify competitors of DNA binding by an ApiAP2 protein, we used AP2-EXP (PF3D7_1466400), which is the only ApiAP2 protein whose AP2 domain is structurally characterized (PDB accession: 3IGM)(36). *In silico* molecular docking was run on AUTODOCK(37) using over ten thousand small molecules from both the Tres Cantos Antimalarial Set (TCAMS)(38) and the Drug Bank(39) against AP2-EXP. Docking hits were prioritized based on the free energy of interaction and proximity to amino acids necessary for DNA contact **(Fig S1A)**. In some cases, the compound that generated a high docking score was not available for purchase, so alternative choices with >0.9 Tanimoto similarity score were identified, computationally docked, and used for further evaluation **(All compounds listed in Figure S1B).** These docking simulations resulted in a final list of nine top scoring predicted competitors of DNA binding by AP2-EXP. Each compound was assigned an alphabetical identifier A-I **(Fig 1A, Fig S2A, B)** for use in this study. We determined that all nine compounds kill asexual *P. falciparum* parasites at micromolar concentrations (range 11-170µM) in a growth inhibition assay **(Fig 1A, Fig S3)**.

**Figure 1.**
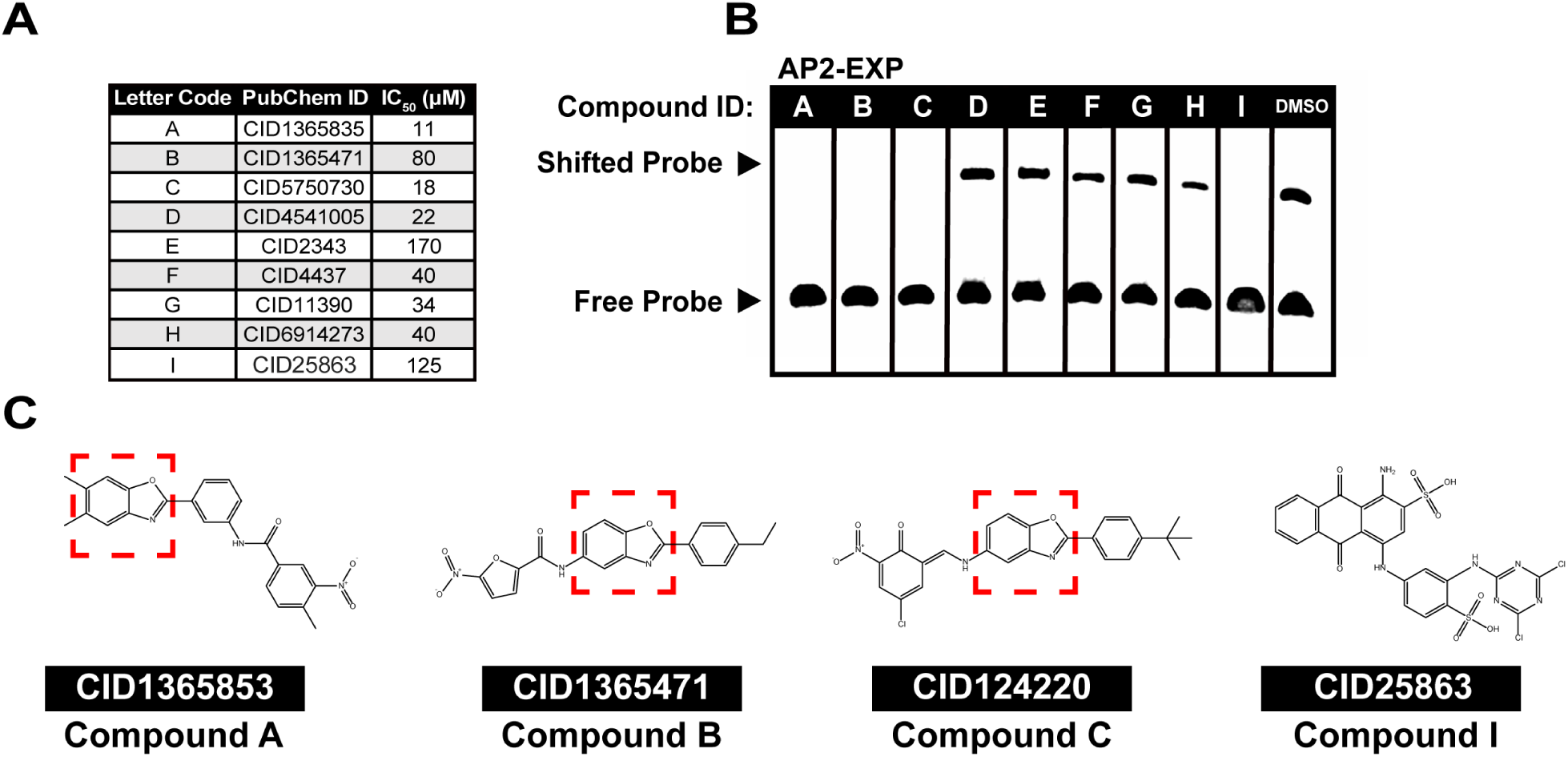
Nine putative competitors of DNA binding by AP2-EXP were identified by an *in-silico* screen and tested *in vitro* and *in vivo*. A) Each compound that was prioritized based on the *in*-*silico* screen was assigned an identifier (A-I) and tested for anti-*Plasmodium* activity in a 48-hour growth inhibition assay against asexual blood stage *P. falciparum*. B) Each putative AP2-EXP competitor was added to an EMSA containing the AP2-EXP AP2 domain. DNA binding competition will result in a loss of the shifted DNA probe. DMSO vehicle was used as a control for normal DNA binding by AP2-EXP. 150 fmoles of DNA probe, 125ng of AP2-EXP, and 300μM of each compound were used for each lane. C) Chemical structures of AP2-EXP competing Compounds A, B, C, and I. Three of the four compounds that compete DNA binding *in vitro* (Compounds A, B, and C) have a benzoxazole core moiety, denoted by a red box.

### Four Compounds Have Activity Against AP2-EXP *in vitro*

The nine predicted competitors of DNA binding by AP2-EXP were tested in a competitive electrophoretic mobility shift assay (EMSA). The minimum mass of AP2-EXP required to visualize DNA binding **(Fig S4A)** was mixed with each compound and incubated prior to adding the DNA oligonucleotide **(Table S1)**. Four of the nine compounds (A, B, C and I) **(Fig 1B)** were able to effectively compete for DNA binding by AP2-EXP *in vitro*. Three of the four AP2 domain competitors (Compounds A, B and C) have a benzoxazole core moiety, while Compound I is made up of planar rings **(Fig 1C)**.

To assess whether DNA binding competition was specific to the AP2-EXP AP2 domain, we repeated competitive EMSAs with three different purified *P. falciparum* AP2 domains: AP2-I Domain 3 (AP2-I D3)(12, 18) **(Fig S4B)**, AP2-HS Domain 1 (AP2-HS D1)(12, 35) **(Fig S4C)**, and PfSIP2 Domain 1 (PfSIP2 D1)(20) **(Fig S4D)**, again using the minimum amount of protein needed to visualize interaction with the DNA oligonucleotide **(Table S1)**. We found that Compounds A, B, C, and I can compete DNA binding by AP2-I **(Fig S5A)**, Compounds B and I compete DNA binding by AP2-HS D1 **(Fig S5B)** and Compounds B, C and I compete DNA binding by PfSIP2 D1 **(Fig S5C)**.

As a control for compound specificity, we performed competitive EMSAs using two non-*Plasmodium* AP2 domain proteins: the *Arabidopsis thaliana* Ethylene Response Factor 1(40) (AtERF1) **(Fig S4E)** and the human High Mobility Group Box (HMGB) domain protein SOX2 **(Fig S4F)**(41). Only Compound I was able to compete AtERF1 **(Fig S6A)**, while SOX2 was competed by Compound A only **(Fig S6B)**. Therefore, Compounds B and C demonstrate specificity for Apicomplexan AP2 domains *in vitro*. (EMSA Results Summarized in **Table S2**).

To eliminate the possibility that ApiAP2 competing compounds interfere non-specifically with DNA binding proteins by intercalating with DNA, we conducted an ethidium bromide exclusion assay(42). Compared to the positive control DRAQ5(42), only Compound I was able to effectively displace ethidium bromide from DNA **(Fig S7)**. Based on these results, we focused on Compounds B and C for further study.

### Structural Analogs of Compound B Differ in DNA Binding Competition Activity

We tested the four closest available analogs to Compound B from the TCAMS based on similarity score(38) (Designated as Compounds B-1, B-2, B-3 and B-4) for competition against AP2-EXP. Compounds B-1 and B-4 compete DNA binding by AP2-EXP **(Fig 2A)** and AP2-I D3 **(Fig S8A)**, while Compounds B-2 and B-3 do not compete DNA binding by either AP2-EXP or AP2-I D3 **(Fig 2A, S8A)**. Compound B-1 is the least effective competitor relative to Compounds B and B-4 **(Fig 2B)**. Non-AP2 domain competitor Compounds B-2 and B-3 each have halogen atoms substituted on the benzene ring. Conversely, Compounds B, B-1, and B-4, have methyl or ethyl groups substituted onto the benzene ring and can compete DNA binding **(Fig 2C)**. Compound B-1 has both a methyl group and a chlorine atom substitution. If the halogen atom substitution decreases DNA binding competition, this mix of substitutions is consistent with its lower efficacy of DNA binding competition relative to Compounds B and B-4. Compounds B, B-1 and B-4 have no difference in DNA intercalation ability measured by ethidium bromide exclusion **(Fig S8B)**. A molecular dynamics simulation of AP2-EXP docked with Compound B and each analogue predicts that Compounds B, B-1, B-2, and B-4 dock stably with AP2-EXP, while non-AP2 competitor Compound B-3 moves away from the protein **(Supplemental Movies 1-5)**. In summary, substituting bulky, electronegative groups onto Compound B can abolish competition for DNA binding with AP2-EXP.

**Figure 2.**
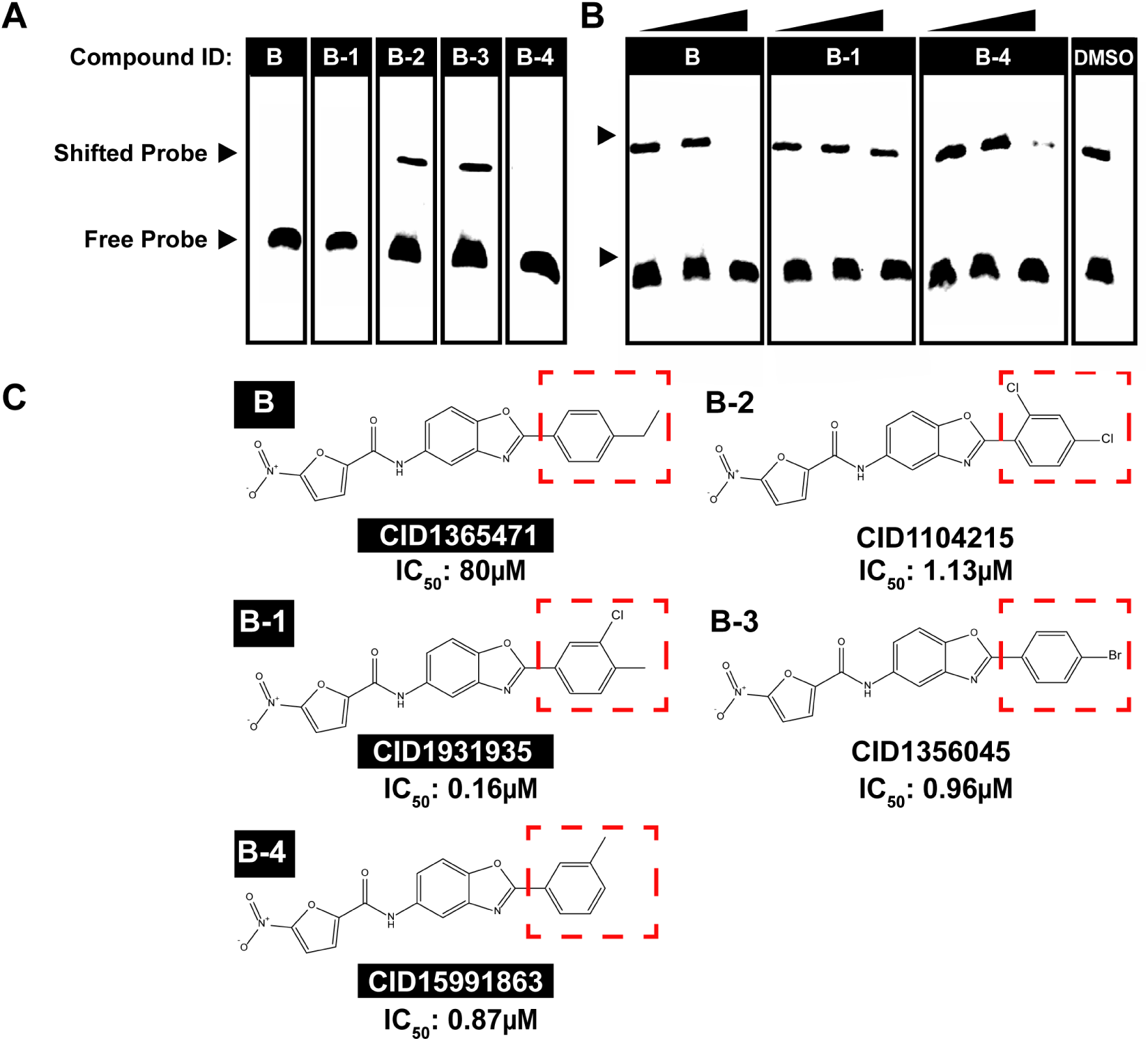
Analogues of Compound B compete AP2-EXP DNA binding with differing efficacy. A) Compounds B, B-1, B-2, B-3, and B-4 were added to an EMSA with AP2-EXP. DNA binding competition will result in the loss of the shifted probe. 40 fmoles of DNA probe, 125ng of AP2-EXP, and 300μM of each compound were used for each lane. B) Compounds B, B-1 and B-4 were titrated at 25, 50, and 150µM into an EMSA with AP2-EXP to determine whether there are differences in DNA binding competition. DMSO vehicle was used as a control for normal DNA binding. 40 fmoles of DNA probe and 125ng of AP2-EXP were used for each lane. The concentration increase for each compound is indicated by the triangle from left to right. C) Chemical structures of Compounds B, B-1, B-2, B-3, and B-4. Compounds that compete DNA binding by AP2-EXP are highlighted in black. Each compound has a different substitution pattern on the right-side benzene ring, denoted by a red box. The IC_50_ against asexual *P. falciparum* of compounds B-1, B-2, B-3, and B-4 measured by Gamo *et al*(38) is indicated at the bottom of each identifier. The IC_50_ of Compound B was determined in this study.

### Compound C Activity Coincides With AP2-EXP Expression

To determine the stage in the IDC at which AP2-EXP is expressed, we used Selection Linked Integration (SLI)(43) to create a parasite line with endogenously GFP tagged AP2-EXP (AP2-EXP::GFP) **(Fig S9A)**. We observed maximum abundance of AP2-EXP::GFP in 30 hpi trophozoites by both western blot and live fluorescence microscopy **(Fig 3A, Fig S10, S11)**. Maximum protein abundance of AP2-EXP in mid-trophozoites was independently confirmed in a parasite line expressing AP2-EXP tagged with 2XHA and the TetR:DOZI mRNA repression aptamer (AP2-EXP::HA) **(Fig S9B, Fig S12)**. AP2-EXP was not detected in maturing Stage III gametocytes **(Fig 3A)**.

**Figure 3.**
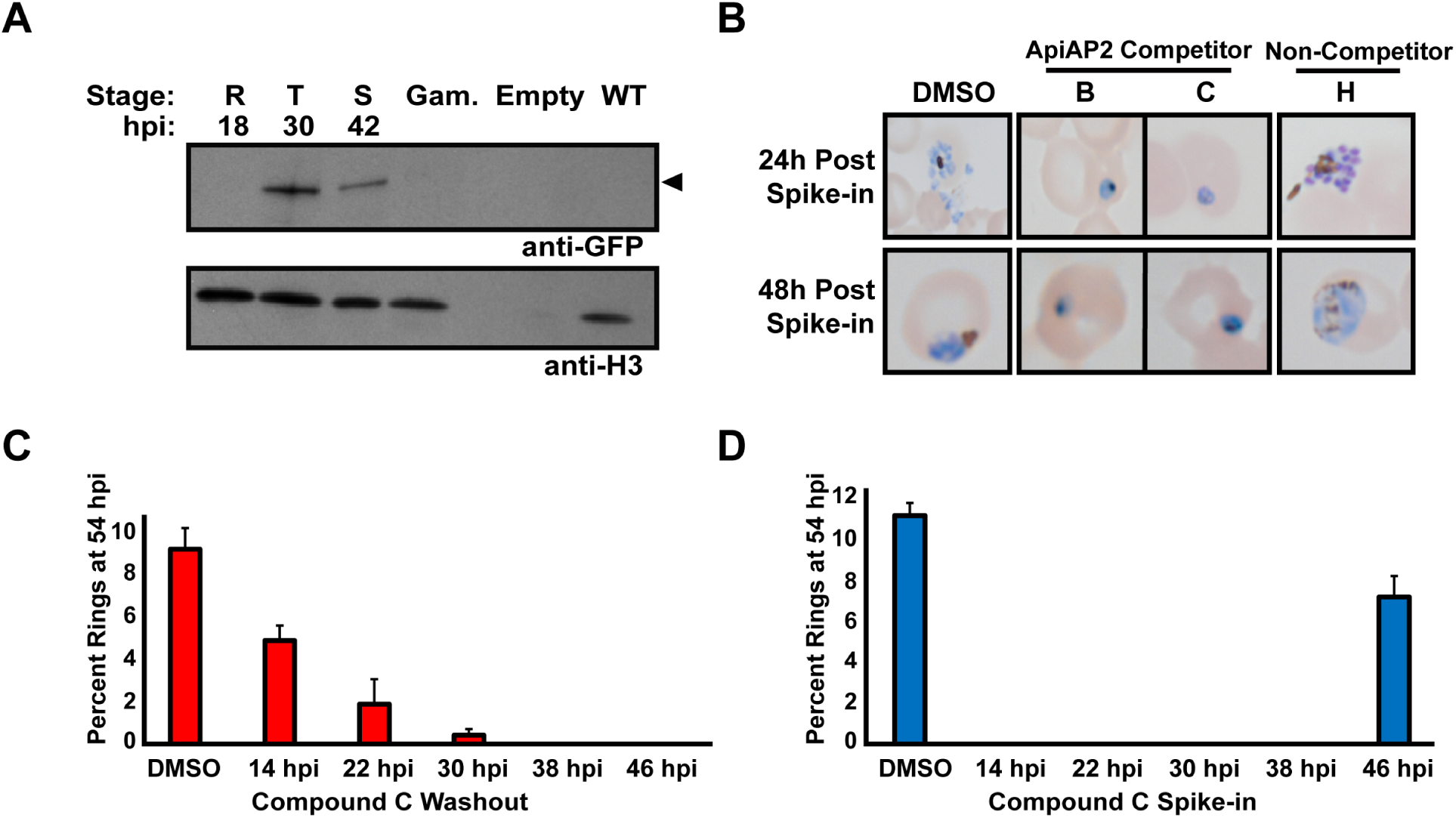
AP2 Domain Competing Compounds B and C affect *P. falciparum* at in the mid trophozoite stage, coinciding with the maximum expression of AP2-EXP. A) AP2-EXP protein expression in the asexual blood stages ring (R), trophozoite (T), and schizont (S), and Stage III gametocytes (Gam), was probed by western blot against AP2-EXP endogenously tagged with GFP (AP2-EXP::GFP). The expected molecular weight of AP2-EXP::GFP (147kDa) is indicated by an arrow. Wild type Pf3D7 protein was used as a negative control. The full length western blot is provided in **Figure S10**. B) Representative Giemsa-stained microscopy images of *P. falciparum* parasites spiked with 40µM Compounds B, C, or H. DMSO vehicle was used as a control for normal progression through the asexual blood stage at 24 and 48 hours post spike-in. C) Highly synchronous asexual blood stage *P. falciparum* parasites were cultured with 40µM Compound C, which was washed out of the media at eight-hour intervals. IDC progression was measured by counting ring stage parasites at 54 hpi. Each assay was done in triplicate. Error bars represent standard deviation. D) Highly synchronous asexual blood stage *P. falciparum* parasites were spiked with 40µM Compound C at eight-hour intervals in the IDC. IDC progression was measured by counting ring stage parasites at 54 hpi Each assay was done in triplicate. Error bars represent standard deviation.

To assess whether Compounds B and C are likely to share a common ApiAP2 target, we determined their parasite killing phenotypes against *P. falciparum* during the IDC. Compound H was used as a control since it does not compete AP2-EXP **(Fig 1B)**. Each compound was spiked (40µM) into synchronous asexual blood stage *P. falciparum* parasites at 24 hpi and parasite morphology was determined at 24 and 48 hours post spike-in **(Fig S13A)**. At 24 hours post spike-in, parasites in the presence of either Compounds B or C had failed to progress past the mid-trophozoite stage and did not morphologically progress any further by 48 hours post spike-in **(Fig 3B)**. In contrast, parasites spiked with Compound H progressed through one IDC before failing to reinvade in the next cycle **(Fig 3B)**. As expected, each compound exposure prevented further parasite growth within 48 hours of spike-in **(Fig S13B)**.

Due to its commercial availability, we continued with Compound C as the lead compound for phenotypic and molecular characterization. To determine the precise timing of Compound C antimalarial activity, synchronous parasites were monitored throughout the 48-hour IDC in the presence of either 40µM Compound C or DMSO control after spiking in Compound C at 6 hpi. Compound C spiked parasites failed to progress beyond the mid-trophozoite stage at 24-30 hours post invasion **(Fig S14)**. To further narrow down the timing of the anti-*Plasmodium* action of Compound C, 40µM Compound C was washed out of a parasite culture at 8-hour intervals beginning at 14 hpi. Growth was rescued, albeit with diminishing effectiveness, until 30 hpi **(Fig 3C)**. Spiking 40µM Compound C into culture at 8-hour intervals beginning at 14 hpi effectively killed the parasite culture at each time point except for 46 hpi. **(Fig 3D)**. Taken together, these results suggest that Compound C maximally inhibits the IDC progression of *Plasmodium* parasites between 30-46 hpi. These results show that the maximum abundance of AP2-EXP at 30 hpi coincides with the timing of action for Compound C.

### Compound C Disrupts the *P. falciparum* Transcriptome with Stage Specificity

Since AP2-EXP is predicted to act as a transcription factor(17, 21) we hypothesized that changes in parasite transcript abundance should occur when AP2-EXP DNA binding is competed by Compound C. We cultured parasites in 12µM (0.66xIC_50_) Compound C or DMSO vehicle and measured RNA abundance at seven time points during the asexual blood stage. As a quality control for normal IDC progression, we found high correlation for a set of “control” genes(44, 45) with an established 48-hour periodicity in the IDC between our DMSO and Compound C samples **(Fig S15A-D)**. Since 12µM Compound C kills fewer than 50% of parasites within one IDC **(Fig S16A, B)**, this demonstrates that there is not an overall perturbation to the IDC transcriptome upon Compound C incubation. The Spearman Correlation between total transcriptome samples deviated most significantly at 30 hpi, (Corr. = −0.16) **(Fig 4A)** as expected from the compound activity measured **(Fig 3C, D, Gene Expression values in Table S3)**.

**Figure 4.**
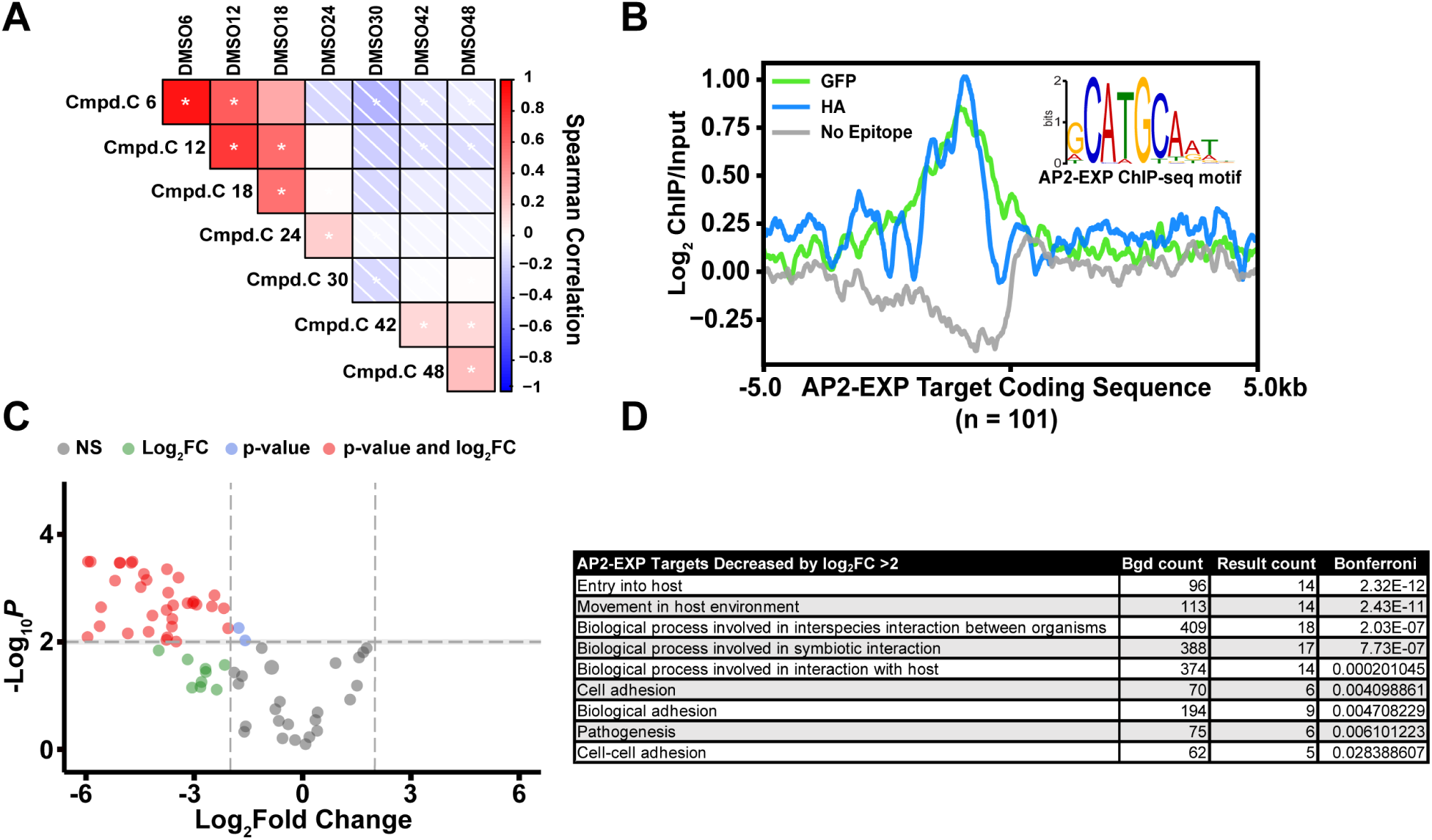
Compound C disrupts the *P. falciparum* transcriptome specifically at 30hpi, with bias towards AP2-EXP targets predicted by ChIP-seq. A) *P. falciparum* parasites were spiked with 12µM (0.66xIC_50_) Compound C or DMSO vehicle control at 10 hpi. Total RNA was harvested at 6, 12, 18, 24, 30, 42, and 48 hpi for quantification by DNA microarray. The full transcriptome Spearman Correlation between Compound C and DMSO control spiked parasites was plotted as a correlogram. A * indicates *p-value* < 0.05. B) Three replicates of AP2-EXP Chip-seq were collected at 30 hpi using two genetically tagged parasite lines (AP2-EXP::GFP and AP2-EXP::HA). A no epitope control was collected using wild type Pf3D7 parasites and anti-GFP antibodies. Log_2_ enrichment of the immunoprecipitate (ChIP) over Input DNA for one replicate of AP2-EXP::GFP, AP2-EXP::HA, and No Epitope Control is plotted relative to the coding sequence start site of AP2-EXP target genes identified in at least 2/3 replicates. CATGA is the most overrepresented DNA motif within the peaks of AP2-EXP occupancy conserved in at least 2/3 ChIP-seq experiments. C) A volcano plot of the changes in abundance for AP2-EXP target genes at 24-30 hpi in the presence of Compound C. 46/93 detected AP2-EXP target transcripts decrease in abundance by log_2_ fold change > 2. D) GO-term analysis of transcripts that are both AP2-EXP targets and decreased in abundance with respect to Compound C (Bonferroni *p-value* cutoff < 0.05).

**Figure 5.**
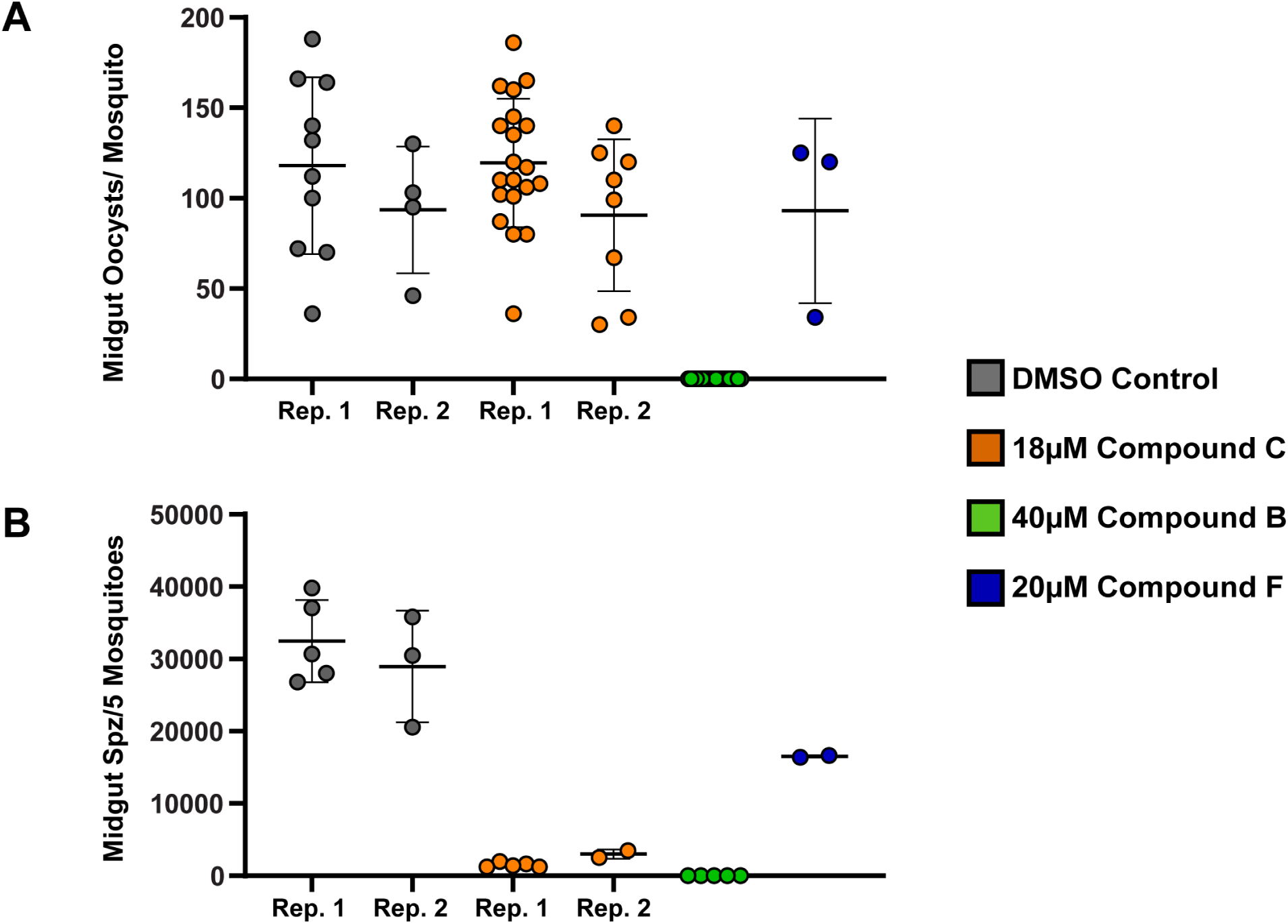
Compounds B and C are active against mosquito stage *Plasmodium berghei* parasites. A) Midgut oocyst counts per mosquito for *P. berghei* infected mosquitoes on day 14 post infection following compound injection on day 10 post infection. Compound identity is indicated in the legend. Rep. 1 and Rep. 2 correspond to biological replicates one and two, respectively. B) Midgut sporozoite counts per five infected mosquitoes on day 14 post infection following compound injection on day 10 post infection. Compound identity is indicated in the legend. Rep. 1 and Rep. 2 correspond to biological replicates one and two, respectively.

Next, we quantified differences in transcript abundance that are unique to the progression from 24 to 30 hpi for Compound C vs. DMSO vehicle control populations using Linear Models for Microarray Data (LIMMA(46)) **(Differentially Expressed genes in Table S3)**. Overall, 463 RNA transcripts are decreased in abundance by log_2_ fold change >2 between 24-30 hpi in the presence of Compound C. Gene ontology (GO) analysis(47) **(Full GO Terms in Table S3)** revealed an enrichment for genes that encode proteins important for the parasite to invade or modify red blood cells among decreased transcripts (e.g., RON3, GAP45, RhopH2, RhopH3, MSP1, MSP6). We independently determined differential transcript abundance over the entire time course using the RNA time course specific software Rnits(48) and found high overlap with our LIMMA analysis **(Overlaps in Table S3)**. Therefore, the maximal perturbation of the transcriptome **(Fig 4A)**, action of Compound C **(Fig 3C, D)**, and maximal abundance of AP2-EXP **(Fig 3A)** all occur at roughly 30 hpi in the IDC.

### AP2-EXP Gene Targets Correlate with Differentially Abundant Transcripts

We used ChIP-seq to determine the genome wide binding occupancy of AP2-EXP. A total of three samples from highly synchronous trophozoites were collected at 30 hpi using both the AP2-EXP::GFP (2 replicates) and AP2-EXP::HA (1 replicate) parasite lines **(Peaks in Table S4)**. As a quality control, we determined that ChIP recovers intact AP2-EXP **(Fig S17A)**, and a co-immunoprecipitation blot confirmed that AP2-EXP interacts with Histone H3 in the nucleus **(Fig S17B)**. In aggregate, AP2-EXP binds 240 genomic loci, corresponding to 101 total target genes **(Genes in Table S4)**. Enrichment of AP2-EXP was well conserved between the GFP and HA tagged parasite lines, as indicated by nearly identical coverage in a metagene plot of AP2-EXP target genes **(Fig 4B)**. A ‘no epitope control’ ChIP done on a wild type Pf3D7 parasite line with the anti-GFP antibody detected only 2 peaks, neither of which overlapped with those of AP2-EXP::GFP or AP2-EXP::HA **(Figure 4B, Table S4)**. AP2-EXP peaks were highly enriched for the known AP2-EXP and PbAP2-Sp DNA sequence motif CATGCA(17, 49) **(Fig 4B, Fig S18A-C)**.

Out of the 463 transcripts decreased in abundance by log_2_ fold change > 2 in the presence of Compound C, 46 are AP2-EXP targets **(Table S3)**. This represents 50% (46/93) of AP2-EXP target genes detected in the RNA time course **(Fig 4C, Figure S19)**. In general, the transcripts with the greatest decrease in abundance are AP2-EXP targets. We found that 7/14 total transcripts decreased in abundance by log_2_ fold change > 5, and 16/33 decreased in abundance by log_2_ fold change > 4, are AP2-EXP targets **(Fig 4C, Table S3)**. AP2-EXP targets that are dysregulated at 24-30 hpi have functions related to red blood cell invasion and host remodeling **(Fig 4D, Table S3)**. To assess whether AP2-EXP DNA binding is impacted by Compound C, parasite cultures were spiked with either 40µM Compound C or DMSO vehicle control at 30hpi and AP2-EXP occupancy was measured by ChIP-quantitative PCR (ChIP-qPCR). AP2-EXP DNA binding decreases at five specific peaks of occupancy (GAP45, SIP2, RON3, RALP, and AMAI) in the presence of Compound C **(Fig S20)**.

We then compared AP2-EXP genomic occupancy with several published datasets in order to further evaluate its function as a sequence specific transcription factor. AP2-EXP occupancy is correlated with a nucleosome depleted region **(Fig S21A, B)**(50) and activating chromatin marks(51, 52) **(Fig S22A, B)**, and is proximal to target gene transcription start sites (TSS)(53) **(Fig S23)**. The majority of target genes increase in abundance starting at 32 hpi **(Fig S24)**(54). AP2-EXP target gene function is enriched for invasion and red blood cell modification, **(Table S3)** as was found for the genes that decrease in abundance in the presence of Compound C **(Fig 4D, Table S3)**. In aggregate, these findings suggest that Compound C inhibits the function of AP2-EXP as a sequence specific transcription factor.

### ApiAP2 Competitor Compounds B and C are Active Against Mosquito Stage *Plasmodium* Parasites

Elucidation of the role of essential *P. falciparum* ApiAP2 proteins in the IDC is challenging due to the inability to knock out asexual blood stage essential genes. We did not recover transgenic parasites after attempting to completely disrupt the coding sequence of AP2-EXP using the targeted gene disruption (pSLI-TGD) system(43) in three independent attempts, suggesting that AP2-EXP is essential for the IDC (not shown). We then attempted to disrupt AP2-EXP protein abundance with the conditional knockdown approaches Knock Sideways(43), TetR:DOZI mRNA repression(55), and *glms* ribozyme mediated cleavage(56). Each of these genetic systems failed to mediate protein knockdown or mislocalization **(Fig S9A, B, Fig S25, Fig S26)**. To account for this limitation, we tested the ApiAP2 competitor Compounds B and C against rodent infectious *Plasmodium berghei* parasites due to the well-characterized genetic phenotype of the AP2-EXP orthologue PbAP2-Sp (89% amino acid identity to the AP2-EXP AP2 domain)(32, 49) as a master regulator of sporogenesis.

We injected the measured IC_50_ **(Figure 1A)** of Compounds B, C, or F into the midgut of *Anopheles* mosquitoes infected with *P. berghei* parasites to determine their effect on *Plasmodium* mosquito stage development **(Fig S27)**. Compound F **(Fig 1B)** was used as a non AP2 domain competing control. Mosquitoes injected with Compounds F or C developed comparable oocyst numbers to the DMSO control **(Fig 6A)** while Compound B prevented the development of midgut oocysts entirely **(Fig 6A)**. Compound C treated mosquitoes developed 10-fold fewer midgut sporozoites than the control **(Fig 6B)**. Therefore, consistent with their hypothesized *in vitro* activity against ApiAP2 proteins, Compounds B and C are strong inhibitors of *P. berghei* sporozoite development *in vivo* (parasite counts in **Table S5)**.

## Discussion

ApiAP2 transcription factors are unique to Apicomplexan parasites due to their plant-like domain architecture(13, 14) and many are essential to asexual blood stage development, making them valuable as potential drug targets. It is therefore desirable to discover chemical scaffolds that can target essential ApiAP2 proteins. Furthermore, chemical inhibition of ApiAP2 proteins may be used in future studies to uncover details about their true biological functions during parasite development. Using a combination of *in silico*, biochemical, and genetic approaches, we identified a series of compounds (Compounds A, B, C, and I) that compete DNA binding by ApiAP2 proteins. Compounds A, B, and C all have the same core benzoxazole moiety but vary in their measured IC_50_, indicating that changes which do not affect ApiAP2 DNA binding competition can alter the potency against parasites. Furthermore, Compounds B and C can inhibit *P. berghei* parasite development without killing mosquitoes, suggesting that ApiAP2 competitor compounds may also be tolerated by the host and may serve as transmission blocking agents(57).

Compounds B and C have no activity against the plant encoded AtERF1 or human encoded SOX2, supporting their selectivity for Apicomplexan AP2 domains. Although the mode of DNA binding is shared between AP2-EXP and AtERF1(40), the specific amino acids that contact DNA are not strictly conserved(36). Lack of activity against AtERF1 may be the result of filtering molecular docking hits based on proximity to DNA base contacting amino acids. The DNA binding competition capacity of Compound B analogs varies *in vitro* based on substitutions to the benzene ring. In a molecular dynamics simulation, Compound B remains stably associated with AP2-EXP, while the non ApiAP2 competitor Compound B-3 does not. This suggests that a steric clash between AP2-EXP and the bromine atom on Compound B-3 is responsible for its lack of DNA binding competition. By extension, this may explain the lower efficacy of DNA binding competition observed for Compounds B-1, B-2, and B-3 compared to B and B-4.

Based on the description of its genome-wide DNA binding sites and transcript dysregulation in the presence of Compound C, new inferences can be made about the biological function of AP2-EXP. This demonstrates the utility of combining chemical and traditional genetics, because previous attempts to genetically characterize AP2-EXP have not provided a complete picture of its function in the asexual blood stage. The *ap2-exp* AP2 domain coding region has previously been shown to be essential for the asexual blood stage by both saturating mutagenesis and targeted deletion attempts(15–17, 21). Conversely, the coding region beyond the AP2 domain was truncated in two studies(15, 21). Transcriptomic profiling of a truncated AP2-EXP parasite line revealed that many transcripts encoding exported proteins were dysregulated(21). Unexpectedly, AP2-EXP target genes identified in this study overlap poorly with the differentially regulated transcripts reported as a result of truncation of AP2-EXP **(Table S4)**(21). There is greater overlap between our study and AP2-EXP ChIP-seq recently described by Shang *et al*(17), with 50/101 target genes conserved **(Table S4)**. This may explain the apparent essentiality of the AP2 domain, while the full-length AP2-EXP has roles that impact a different subset of non-essential genes. A comparison of transcripts differentially abundant in the presence of Compound C to the DNA binding occupancy of AP2-I(18) revealed that 54/85 AP2-I target genes are decreased in abundance by log_2_ fold change >2 at 24-30 hpi. Interestingly, 22/55 AP2-I target genes that decrease in abundance when parasites are exposed to Compound C are also AP2-EXP targets, implying the potential for co-regulation of certain gene subsets. This overlap is consistent with our EMSA results and corroborates the hypothesis that Compound C competes both AP2-EXP and AP2-I *in vivo*. Only 3/22 PfSIP2 target genes are decreased in abundance in the presence of Compound C **(ApiAP2 Target Gene Results Summarized in Table S3)**. This may reflect the cryptic relationship between PfSIP2 DNA binding and transcriptional control, as was noted when PfSIP2 was originally characterized as playing a role in genome integrity and heterochromatin formation(20). Several ApiAP2 proteins have been implicated in oocyst and sporozoite development in a *P. berghei* ApiAP2 knockout screen(24). Since Compounds B and C both inhibit mosquito stage development of *P. berghei* with different phenotypes, it is possible that they target different sets of ApiAP2 proteins *in vivo*. Since PbAP2-Sp is required for sporogony, Compounds B and C should minimally inhibit sporozoite development, which is consistent with our results. Overall, these data support a potential for multi-AP2 domain competition by Compounds B and C *in vivo*. Due to their inclusion in the TCAMS, Compounds B, B-1, B-2, B-3, and B-4 have all been tested for activity against the human HEPG2 cell line(38). ApiAP2 competitor Compounds B and B-4 inhibit HEPG2 cell growth by just 4 and 8%, respectively(38). Therefore, either drug may potentially be prioritized for further development based on selectivity for *Plasmodium* parasites.

AP2-EXP remains the only ApiAP2 protein for which the structure has been solved. Surprisingly, over a decade later, the field still lacks a clear understanding of the biological role of AP2-EXP or insights into the druggability of the AP2 domain. Our study has enabled us to make inferences about the role of AP2-EXP and set a proof of principle for targeting the highly unique Apicomplexan AP2 DNA binding proteins as a new antimalarial strategy.

## Materials and Methods

### Parasite lines

Parasites were grown at 37°C, 5% O2, 7% CO2 using RPMI1640 media supplemented with hypoxanthine and .5% Albumax II (Thermo). The parasite lines used in this study were AP2-EXP::GFP, AP2-EXP::HA, AP2-EXP::*glms*, and wild Type Pf3D7 (Malaria Research and Reference Reagents Repository). The human biological samples were sourced ethically and their research use was in accord with the terms of the informed consents under an IRB/EC approved protocol.

To create AP2-EXP::GFP the C-Terminal coding region of AP2-EXP was cloned **(Table S1)** into the plasmid pSLI::2xFKBP(43) for endogenous tagging by single homologous recombination **(Fig S9A)**. Transgenic parasites with the correct C-Terminal tag were further transfected with pLyn-FRB-mCherry(43) for inducible mislocalization using the same method. Parasites were maintained in media with 2.5nM WR99210. All ChIP and western blot experiments using AP2-EXP::GFP except **Figure S26C** were collected using AP2-EXP::GFP without the pLyn mislocalization plasmid.

To create AP2-EXP::HA we used the pSN054 vector system(58). The right homology region (RHR) was amplified by PCR **(Table S1)**, and the recodonized left homology region (LHR) was synthesized using the BioXP™ 3200 System. Single guide RNA fragments synthesized using the BioXP™ 3200 System **(Table S1)** were cloned into the linearized pSN054 donor vector(58) The parental parasite line used for transfection expresses Cas9 and T7 RNA polymerase(59) **(Fig S9B)**. Cell cultures were maintained in 500nM anhydrotetracycline (aTc, Sigma-Aldrich 37919) and 2.5 mg/mL of Blasticidin S.

To create AP2-EXP::*glms*::HA, the C-terminal coding region of AP2-EXP **(Table S1)** was cloned into the plasmid pSLI::3xHA::*glms*(60) **(Figure S25)**. Parasite cultures were maintained in media with 2.5nM WR99210.

All *P. falciparum* transfection were performed as described(43, 61). All parasite strains were cloned by limiting dilution and genotyped using PCR. AP2-EXP::GFP and AP2-EXP::HA were also genotyped using whole genome sequencing **(NCBI SRA: PRJNA818769)**.

### Genomic DNA Isolation

Parasite cultures were lysed with .1% Saponin in 1xPBS and collected by centrifugation at 1500 RPM, then resuspended in 1xPBS. Genomic DNA was then isolated using the Qiagen DNeasy nucleic acid isolation kit according to the manufacturer’s instructions.

### AP2-EXP Knockdown Assays

For AP2-EXP::GFP + pLyn, synchronous parasites were split into two populations. 250nM Rapalog was added to one parasite group for 48 hours as described(43) while the second group was used as a control. Addition of Rapalog did not affect parasite growth (not shown). After 48 hours parasite protein was collected in nuclear and cytosolic fractions and AP2-EXP::GFP, Histone H3, and Pf Aldolase were detected by western blot.

For AP2-EXP::HA parasites were grown routinely in the presence of 500nM aTc. aTc was washed out of one group, while the second group was maintained with aTc as a control for 120 hours before protein harvest. Parasite growth was not affected by removal of aTc (not shown). AP2-EXP::HA and Histone H3 were detected by western blot.

For AP2-EXP::*glms*::HA, synchronous parasites were split into two populations. 5mM glucosamine was added to one parasite group for 72 hours as described(56) while the second group was used as a control. Addition of glucosamine did not affect parasite growth (not shown). After 72 hours parasite protein was harvested for detection of AP2-EXP::*glms*::HA and Histone H3 by Western blot.

### *In silico* docking screen

Three dimensional structures of every available molecule from the Tres Cantos Antimalarial Set(38) and the DrugBank(39) were created using BALLOON and the Merck Molecular Force Field. The crystal structure of AP2-EXP (PDB:3IGM) was modeled using AutoDockTools(62) and a molecular docking screen was run using AutoDock(37). Details about the preparation of ligands, the protein macromolecule, and the docking screen criteria are provided in the SI text.

### IC_50_ Determination

The IC_50_ of each putative ApiAP2 competing compound was determined using a 48-hour growth inhibition assay as described(63). Parasites were seeded at .5% parasitemia and 4% hematocrit on a 96 well plate. Drug or DMSO vehicle control was added in triplicate to each well and parasites were incubated in standard culture conditions for 48 hours. Growth values were normalized to the vehicle control. IC_50_ values were calculated and growth inhibition curves were plotted using GraphPad Prism.

### Recombinant Protein Expression

Recombinant AP2 domains AP2-EXP, AP2-I D3, AP2-HS D1, and PfSIP2 D1 were overexpressed and purified from BL21 PlysS *E. coli* as described previously(12). Recombinant AtERF1 was purified using the same method after cloning the AtERF1 AP2 domain(40) into the pGex4t-1 overexpression vector **(Table S1)** Recombinant protein was quantified using Braford reagent (Pierce). Input, flowthrough, and eluate fractions were analyzed by SDS-PAGE to ensure recovery of the full-length recombinant protein. Recombinant full-length SOX2 protein was purchased from Abcam (ab169843).

### Electrophoretic Mobility Shift Assays

EMSAs were performed using the Thermo Light Shift EMSA kit with recommended buffer components. Gel shifts were visualized using the Light Shift detection reagents and imaged using a BioRad Chemiluminescence imager. Additional details are provided in the SI text.

### Ethidium Bromide Exclusion Assay

10µM of double stranded DNA (sequence TGCATGCA, purchased from IDT) in .01mM EDTA, 9.4mM NaCl, 2mM HEPES buffer pH 7.9 was incubated for 10 minutes in the presence of 100nM ethidium bromide. Fluorescence was measured at excitation/emission 546nm/595nm for each well. Following the baseline reading, each putative ApiAP2 competing compound was added. DRAQ5 nucleic acid dye was used as a positive control for knockdown of ethidium bromide fluorescence by DNA intercalation. Ethidium bromide exclusion assays were performed in technical triplicate using a 96 well plate.

### Molecular Dynamics Simulations

Five Compounds (B, B-1, B-2, B-3, and B-4) were simulated using AMBER(64) for 100ns of interaction with the AP2-EXP AP2 domain. The initial position of each compound used was identical to the predicted docking conformation of Compound B **(Figure S2A)**. Additional details are provided in the SI text.

### AP2 Domain Competitor Phenotyping Assays

For Compound B, C, and H spike in phenotyping **(Fig 3B)**, highly synchronous Pf3D7 wild type parasites were spiked with 40µM of each drug at 24 hpi. An equivalent volume of DMSO was used as a control. Parasites were morphologically assessed by Giemsa staining at 24 and 48 hours post spike in (48 and 72 hpi).

For continuous Compound C exposure phenotyping **(Fig S14)**, highly synchronous Pf3D7 wild type parasites were spiked with 40µM Compound C or equivalent DMSO vehicle control at 6 hpi. Parasite progression was then monitored through the IDC by Giemsa staining at 6-hour intervals.

For Compound C spike in and washout assays **(Fig 3C, D)**, 40µM of Compound C was spiked into highly synchronous Pf3D7 wild type parasites at 8-hour intervals starting at 14 hpi. For Compound C washout assays, 40µM of Compound C was washed out of highly synchronous culture at 8-hour intervals starting at 14 hpi. For each assay an equivalent volume of DMSO vehicle was used as a control for normal growth.

40µM of compound was used for all phenotyping assays because it was the maximum concentration achievable in culture after resuspending each compound in 100% DMSO. Higher concentrations of compound caused parasite mortality due to the high volume of DMSO required (>1% volume/volume). All assays were performed in triplicate and parasitemia was counted by Giemsa-stained slides. Error bars represent the standard deviation of the mean.

### Preparation of RNA for DNA Microarray

DNA microarrays were prepared using the protocol described in(44). After collecting a control timepoint at 6 hpi, parasites were cultured with either 12µM Compound C (0.66xIC_50_) or DMSO vehicle control. Parasite RNA was then harvested at 12, 18, 24, 30, 42, and 48 hpi. cDNA was synthesized using SuperScript II, hybridized onto Agilent DNA microarrays(44), and scanned using an Axon 4200A Scanner. Agilent Feature Extraction Software version 11.0.1.1 with the protocol GE2-v5_95_Feb07_nospikein was used to extract signal intensities. Raw signal intensities for all microarrays are available in **Table S6**.

### Analysis of DNA Microarray**s**

Microarray data were extracted and normalized using the R packages LIMMA(46) and Rnits(48). LIMMA was used to analyze differential transcript abundance specific to the 24-30 hpi time points, while Rnits was used to assess differential transcript abundance across the entire IDC. Additional details are provided in the SI text.

### ChIP Sample Preparation

ChIP-seq was performed as described in(19). Highly synchronous parasites were grown up to 5-10% parasitemia and AP2-EXP ChIP was performed at 30 hpi for all replicates. Immunoprecipitation was performed overnight with .5mg/mL 3F10 anti-HA (Sigma) or .1mg/mL Ab290 (Abcam) anti-GFP.

### DNA Library Preparation

ChIP-seq DNA libraries were prepared as described in(19) using the NEBNext II DNA library kit (New England Biolabs) according to the manufacturer’s instructions. Quality was assessed using an Agilent 2100 Bioanalyzer or TapeStation. Libraries were sequenced using a HiSeq 2500 (Replicate GFP1) or NextSeq 550 (Replicates GFP2 and HA3) Illumina sequencer. AMPure XP beads (Beckmann Coulter) were used to size select and purify DNA between NEBNext II library preparation steps. Whole genome sequencing DNA libraries were prepared using the Illumina Tru-Seq PCR free DNA library kit and sequenced using a HiSeq 2500.

### qPCR

ChIP-qPCR samples were collected from 30-35 hpi trophozoites following two hours of Compound C (40μM) or DMSO vehicle control spike in. Primer pairs to be used for ChIP-qPCR were first evaluated to check for 80-110% efficiency using sonicated genomic DNA. RT-qPCR was carried out using Sybr Green Polymerase master mix (Thermo) with the specified primer concentration **(Table S1)**. The Ct was calculated using SDSv1.4 (Applied Biosystems) software, averaged over technical triplicate. The percent of input per immunoprecipitated DNA fraction was calculated using the delta Ct method. Each assay was performed in biological triplicate, with the exception of the *ama1* primer pair, where n =2. Error bars represent standard deviation of the mean. Data was obtained using an Applied Biosystems 7300 Real-Time PCR Machine.

### ChIP-seq data analysis

ChIP-seq reads were mapped to generate bam and bigwig coverage files as described(19). Peaks of occupancy were called using MACS2(65) and peak intervals were analyzed for overlap and gene proximity using BedTools(66). DeepTools(67) and cegr-tools: https://github.com/seqcode/cegr-tools were used to compare AP2-EXP genomic occupancy to previously published datasets. BedTools(66) was used to compare AP2-EXP peaks determined in this study to AP2-EXP occupancy determined in Shang *et al*(17). Additional details are available in the SI text.

### Western blot

Full parasite protein western blot samples were collected by lysing RBCs with .1% saponin and boiling protein in Loading Buffer (50mM Tris-Cl pH 8.0, 20% SDS, 1% Bromophenol Blue). Fractionated parasite protein was prepared as described in(68). Blots were performed as described(19). Primary antibodies used were: 1/1000 rat ant-HA (Roche 3F10), 1/1000 mouse anti-GFP (Roche), 1/3000 rabbit anti-aldolase conjugated to HRP (Abcam ab38905), or 1/3000 mouse anti-H3 (Abcam ab10799). Secondary antibody concentrations used were 1/3000 goat anti-rat HRP conjugate (Millipore), 1/3000 goat anti-mouse HRP conjugate, or 1/10,000 (Pierce) goat anti-rabbit HRP conjugate (Millipore). ECL reagent (Pierce) was used to detect HRP signal. Blots were exposed to autoradiography film (VWR) and visualized using an autoradiography developer.

### Protein Pulldown

Parasite nuclear protein was isolated as described(68). Following the final centrifugation step, the supernatant was collected and diluted by 1:3 in Dilution Buffer (30% glycerol, 20mM HEPES pH 7.8). GFP tagged AP2-EXP was pulled down using Chromotek anti-GFP or mock immunoprecipitated using Chromotek negative control magnetic beads. Beads were washed twice in Wash Buffer prior to use (20mM HEPES pH 7.4, 250mM NaCl, 1mM EDTA, 1mM TCEP, .05% NP-40). Protein and beads were incubated together for 1 hour at 4°C. The beads were then washed twice with Wash Buffer and bound protein was collected in Loading Buffer by boiling at 95°C for 10 minutes.

### Fluorescent Microscopy

Samples were prepared by incubating packed infected red blood cells with DRAQ5 dye (Thermo) for 15 minutes. Parasites were then washed in 1xPBS to remove excess dye and immediately placed on a glass slide for imaging. Fluorescent microscopy images were acquired using an Olympus Bx61 fluorescent microscope. All images were processed using SlideBook 5.0.

### Mosquito Stage *P. berghei* AP2 Competitor Growth Assay

A female Swiss Webster mouse was inoculated with Plasmodium berghei ANKA 2.34 from frozen stock. Once the parasitemia reached 15%, the blood was harvested by heart puncture, washed twice with 1xPBS and resuspended to 10 mL in 1xPBS. 5 female Swiss Webster mice were infected with 500µL of the resuspended blood. 3-4 days after blood passage, exflagellation of the *P. berghei* gametocytes was assayed. Briefly, a drop of tail vein blood was incubated in RPMI 1640 (Invitrogen) containing 1µM xanthurenic acid (Sigma) for ∼12-15 minutes. The mixture was added to a slide and observed under a light microscope at 40x. Ten or more fields were observed. Mice with 0.3-0.7 exflagellations per field were anesthetized and fed on 3-day post emergence *Anopheles stephensi* mosquitoes. Mosquitoes were maintained at 19°C and fed 10% sucrose. Bloodfed mosquitoes were separated 30 hours post-bloodfeeding. 10 days post-bloodfeeding, 5-10 mosquitoes were dissected and oocysts were counted by mercurochrome staining and light microscopy to ensure *P. berghei* oocyst development. The remaining mosquitoes were injected by standard mouth pipette technique with different small molecule inhibitors or a mixture of PBS-DMSO for control. The surviving injected mosquitoes were dissected 14 days post-bloodfeeding. Oocyst numbers were counted as before. Pooled groups of midguts were ground using a pestle, centrifuged at 7,000 rpm for 5 minutes and resuspended in 20µL 1xPBS to release developing mid-gut sporozoites. Sporozoites were counted on a haemocytometer. All *in vivo* studies were conducted in accordance with the GSK Policy on the Care, Welfare and Treatment of Laboratory Animals and were reviewed by the ethical review process at the institution where the work was performed.

## Data Availability

All sequencing data for AP2-EXP ChIP-seq and whole genome sequencing for transgenic parasite lines AP2-EXP::GFP and AP2-EXP::HA have been deposited in the SRA under **PRJNA818769**.

Raw microarray data for the Compound C and DMSO control RNA time course is available in **Table S6**.

## Author Contributions

T.J.R., E.K.D.S., K.S.S., N.D., C.F.A.P., G.P., M.J.L., C.D.O., J.C.N. and M.L. designed experiments. T.J.R., E.K.D.S., V.C., K.S.S., N.D., G.J., C.F.A.P., and I.K. performed the experiments. T.J.R. and N.D. analyzed the data. T.J.R. generated figures. T.J.R. and M.L. wrote the manuscript in collaboration with the other authors

## Acknowledgements

This work was funded through NIH/NIAID R01AI076276 (M.L.), R01AI125565 (M.L.), and with support from the Center for Quantitative Biology (P50 GM071508) (M.L.). T.J.R. was supported by NIH T32 Predoctoral Training Grant (5T32GM125592-01) awarded to the Center for Eukaryotic Gene Regulation (CEGR) at The Pennsylvania State University. G.A.J. is a recipient of the Sir Keith Murdoch Fellowship from the American Australian Association and a Postdoctoral Research Grant from the American Heart Association (16POST26420067).

## SI Text

### Extended Methods

#### *In Silico* Docking Screen

The Tres Cantos Antimalarial Set of 13533 small molecules was retrieved from the supplementary material of Gamo *et al*(1) and downloaded in Spatial Data File (SDF) format from BatchEntrez. 4603 small molecule structures were downloaded from DrugBank(2) version 2.5 in SDF format. Using the SDF library, a 3-dimensional MOL2 library was created with the use of BALLOON and the Merck Molecular Force Field. BALLOON was used to calculate up to 20 conformations using a genetic algorithm and to select the conformation with the lowest energy of conformation. The 3-dimensional MOL2 library was then prepared for AutoDock using the python script “ligand_prepare4.py” from AutoDockTools(3). The prepared ligands were then saved in PDBQT file format. The python script “ligand_prepare4.py” automatically defines the torsions of the molecule.

The homodimerized AP2-EXP AP2 domain was downloaded from the solved crystal structure (PDB ID 3IGM). AutoDockTools was used to remove water molecules, add hydrogens, and calculate charges. Residues Arg-88 and Asn-118 were found to be flexible using FlexPred and set as flexible residues for docking. The flexible residues were saved as flexible residues in a flexible residue file and the remaining molecule was saved as rigid residues in a rigid residue file as specified in the AutoDock manual(3). The rigid residues file was used to pre-calculate an energy grid of the macromolecule using autogrid4(3). A set of 14 grid map types S, Cl, F,A, Br, N, P, OA, SA, C, I, HD, Ca, NA were calculated for the macromolecule using AutoGrid(3).

From the prepared ligands a docking set was prepared for AutoDock with the macromolecule and pre-calculated energy grids. For each docking set a docking parameter file was prepared, with the Lamarckian Genetic Algorithm (LGA) specified as the search algorithm for AutoDock. To allow for high throughput parallel processing PERL was used on top of AutoDock to manage the dockings. The algorithm was initially set to run with a maximum of 25000 energy evaluations and 20 repeats. Since the calculations are computationally intensive, this setting was used as initial screening of the ligands, to identify a subset of ligands for a more thorough evaluation(3). The 1000 best hits from the GSK compound evaluation and the DrugBank compound evaluation were selected for a more thorough evaluation. The docking parameters were reconfigured to 250000 energy evaluations and 100 repeats and docking repeated. The docking results were examined with PERL, to provide an automated approach to the interpretation of the docking results. The docking results were evaluated using two different approaches. The top candidates for competition of DNA were selected using a PERL script to filter based on having a geometric center within 10 Å of the location of sense-strand DNA binding. Hits were then filtered based on having a predicted free energy of interaction < −5 kJ/Mol.

#### Electrophoretic Mobility Shift Assay

EMSAs were run in DNA binding buffer (10mM Tris pH 7.5, 50mM KCl, 1mM DTT, 6mM MgCl2, 60ng/µL Poly DiDC, 65ng BSA). Recombinant proteins were titrated to empirically determine the minimum mass required for DNA binding and this mass **(Fig S4)** was used for each gel-shift unless otherwise specified. PAGE purified DNA probes with a 5’ biotin ligated on the forward DNA strand, along with an unlabeled complementary stand, were purchased from IDT **(Table S1)**. DNA probes were double stranded by heating to 95°C, followed by stepwise cooling in Annealing Buffer (10mMTtris-Cl pH7.5, 1mM EDTA, 10mM NaCl). Each recombinant protein was incubated in DNA binding mixture plus competitor compound for 15 minutes prior to addition of the cognate double stranded DNA oligonucleotide. Protein, competitor, and DNA oligonucleotide were incubated together for an additional 5 minutes. The mixtures were separated on a .5x TBE polyacrylamide gel, transferred to a nylon membrane (Amersham) at 50 Volts for 30 minutes, and probed using the Light Shift nucleic acid detection module (Thermo) according to the manufacturer’s protocol. All gels were imaged using a Bio Rad chemiluminescence imager.

#### Molecular Dynamics Simulations

Five compounds were prepared for molecular dynamics (MD) simulation: B, B-1, B-2, B-3, and B-4. In preparation for MD, parameters for the compounds were obtained using Antechamber(4–6) with the Generalized Amber Force Field. Starting conformations for each compound bound to AP2-EXP were based on predictions from docking. All complexes were prepared using the tleap module of AmberTools(7) with the protein.ff14SB forcefield(8). Each complex was solvated in an octahedral box of TIP3P water with a 10 -Å buffer around the protein complex. Na^+^ and Cl^-^ ions were added to neutralize the protein and achieve physiological conditions. All MD minimizations and simulations were performed using Amber with GPU acceleration(9, 10). First, complexes minimized with 5000 steps each of steepest decent and conjugate gradient minimization with 500 kcal/mol·Å^2^ restraints on all complex atoms. Restraints were reduced to 100 kcal/mol·Å^2^ and the minimization protocol was repeated. Restraints were then retained only on the compound for a final minimization step. Following minimization, all complexes were heated from 0 to 300 K using a 100-ps run with constant volume periodic boundaries and 10 kcal/mol·Å^2^ restraints on all protein and compound atoms. To equilibrate complexes, 10 ns of MD was performed first with 10 kcal/mol·Å^2^, then with 1 kcal/mol·Å^2^ restraints on protein and compound atoms using the NPT ensemble. With kcal/mol·Å^2^ restraints retained on complexes, 500 ns production simulations were performed. A 2-fs timestep was used and all bonds between heavy atoms and hydrogens were fixed with the SHAKE algorithm(11). A cut-off distance of 10 Å was used to evaluate long-range electrostatics with Particle Mesh Ewald (PME) and for van der Waals forces. The ‘strip’ and ‘trajout’ commands of the CPPTRAJ module(12) were used to remove solvent atoms and extract 50,000 evenly spaced frames from each simulation for analysis.

#### Analysis of DNA Microarrays

LIMMA(13) was used to normalize and extract signal intensities. Arrays were normalized using robust splines normalization and within arrays parameters selected. Average expression values were calculated per gene by averaging the log_2_ Cy5 (cDNA)/Cy3 (Reference Pool) signal intensity across all probes. Correlation plots for the Compound C vs. DMSO control total transcriptome and control genes were made using the log_2_ signal intensities for all detected genes using the R package CorrPlot downloaded from: https://github.com/taiyun/corrplot. The LIMMA eBayes function was used on average gene abundance data to determine changes in transcript abundance between 24 and 30hpi that occur differentially in DMSO control vs. Compound C dosed parasites. Volcano plots were made with LIMMA eBayes data using the Enhanced Volcano package in R downloaded from: https://github.com/kevinblighe/EnhancedVolcano. The Rnits(14) R package was used to model differences in gene expression across the entire time course between the DMSO control and Compound C spike in parasites. Rnits was run using the parameters: center genes, normalize by intensity, and background filter probes. The Rnits model for differential expression was fit at the gene level. All data from LIMMA, eBayes, and Rnits is provided in **Table S3**. Heat maps for control genes and AP2-EXP target genes were made using Java Treeview(15) after means centering and log transformation of transcript abundance values with Cluster3.0(16).

#### ChIP-seq Data Analysis

First, reads in each DNA library were trimmed to remove low quality base calls and Illumina adaptor sequences using Trimmomatic(17) with a quality score cutoff of 20. FastQC(18) was used to assess the quality of each DNA library following this step. Reads were then mapped to the *Plasmodium falciparum* genome version 36 downloaded from PlasmoDB using BWA-Mem(19). Multiply mapped reads were filtered out using Samtools(20). Filtered and mapped Input and immunoprecipitate bam files were used for peak calling by MACS2(21) with the following parameters: effective genome size 20000000, No Model, q 0.01. Peaks called in each ChIP-seq replicate were overlapped using Bedtools(22) Intersect, and peaks that occur in a minimum of 2/3 replicates were used for downstream analysis. For data visualization using IGV, .bam files were converted to log_2_ Immunoprecipitate fraction/Input fraction bigwigs using DeepTools(23) BamCompare. Bedtools(22) ClosestBed was used to correlate regions called as peaks of occupancy by MACS2 to the closest gene. Results were filtered based on 1.5kb proximity to the MACS2 peak and strandedness of the putative target gene. For comparison of ChIP-seq library coverages to Transcription Start Sites and Start Codons, DeepTools PlotHeatmap was used. Matrix files underlying PlotHeatmap were created using DeepTools ComputeMatrix. A four-color plot of conserved AP2-EXP DNA binding sites was made using the program cegr-tools four color plot downloaded from https://github.com/seqcode/cegr-tools/tree/master/src/org/seqcode/cegrtools. DNA motifs enriched in AP2-EXP ChIP-seq peaks were determined using the DREME Suite(24). DNA motifs were sorted from highest to lowest level of conservation to the aggregate AP2-EXP DNA motif using FIMO(25).

For comparison of AP2-EXP peaks with existing post-translational histone marks, chromatin post translational modification datasets(26, 27) were downloaded from https://github.com/Daread/plasmodiumExprPrediction(28). Chromatin reader and nucleosome occupancy datasets(29–31) were downloaded from NCBI. ChIP-seq data for BDP1 and HP1 was processed as described above. Nucleosome occupancy data was downloaded from NCBI and processed as described above, with the exception that bam files were turned into bigwig using BamtoBigWig with the --mnase parameter selected. DeepTools ComputeMatrix was used to format the data. DeepTools PlotHeatmap was used to plot coverage of each dataset within 10kb of AP2-EXP peaks. For negative control coverage plots, Bedtools(22) ShuffleBed was used to create random genomic intervals on the same chromosome and the same length as AP2-EXP peaks of occupancy.

For comparison of AP2-EXP peaks identified by Shang *et al*(32) to this study, AP2-EXP peaks of occupancy in the trophozoite stage were downloaded from https://www.ncbi.nlm.nih.gov/geo/query/acc.cgi?acc=GSE184658. Peak summits from each replicate were expanded to 200 nucleotide windows using Bedtools SlopBed(22). Conserved peaks in 2/2 replicates were identified using Bedtools intersect(22), then AP2-EXP peaks from this study and Shang *et al* were overlapped using Bedtools Intersect(22).

---

**Figure S1.**
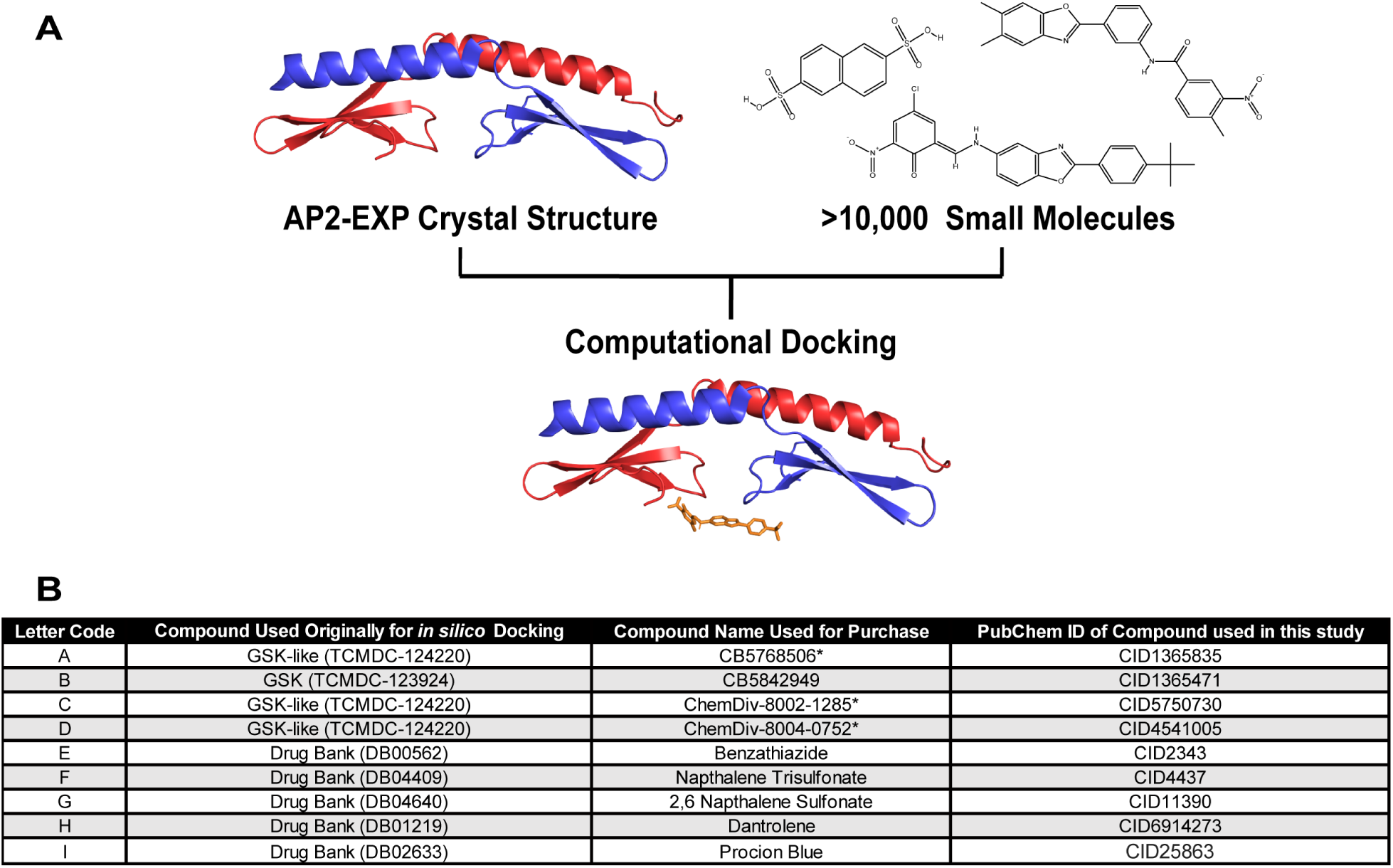
Putative AP2-EXP competitors were identified using computational docking. A) The crystal structure of AP2-EXP (PDB:3IGM)(33) was used as a template to computationally dock thousands of small molecules *in silico* using AutoDock. Results were filtered for compounds that dock within 10 Angstroms of DNA binding residues with a free energy less than −5kJ/mol. Compounds matching these criteria were sourced and used for further testing. B) Seven compounds were identified as putative ApiAP2 competitors in an *in-silico* screen (Column 2). Five of these were available for direct purchase (Column 3). For the two remaining compounds, four alternate choices with a Tanimoto similarity score of .9 or greater were purchased (denoted by an asterisk in Column 3). The PubChem ID used to purchase each compound in this study is listed in Column 4.

**Figure S2.**
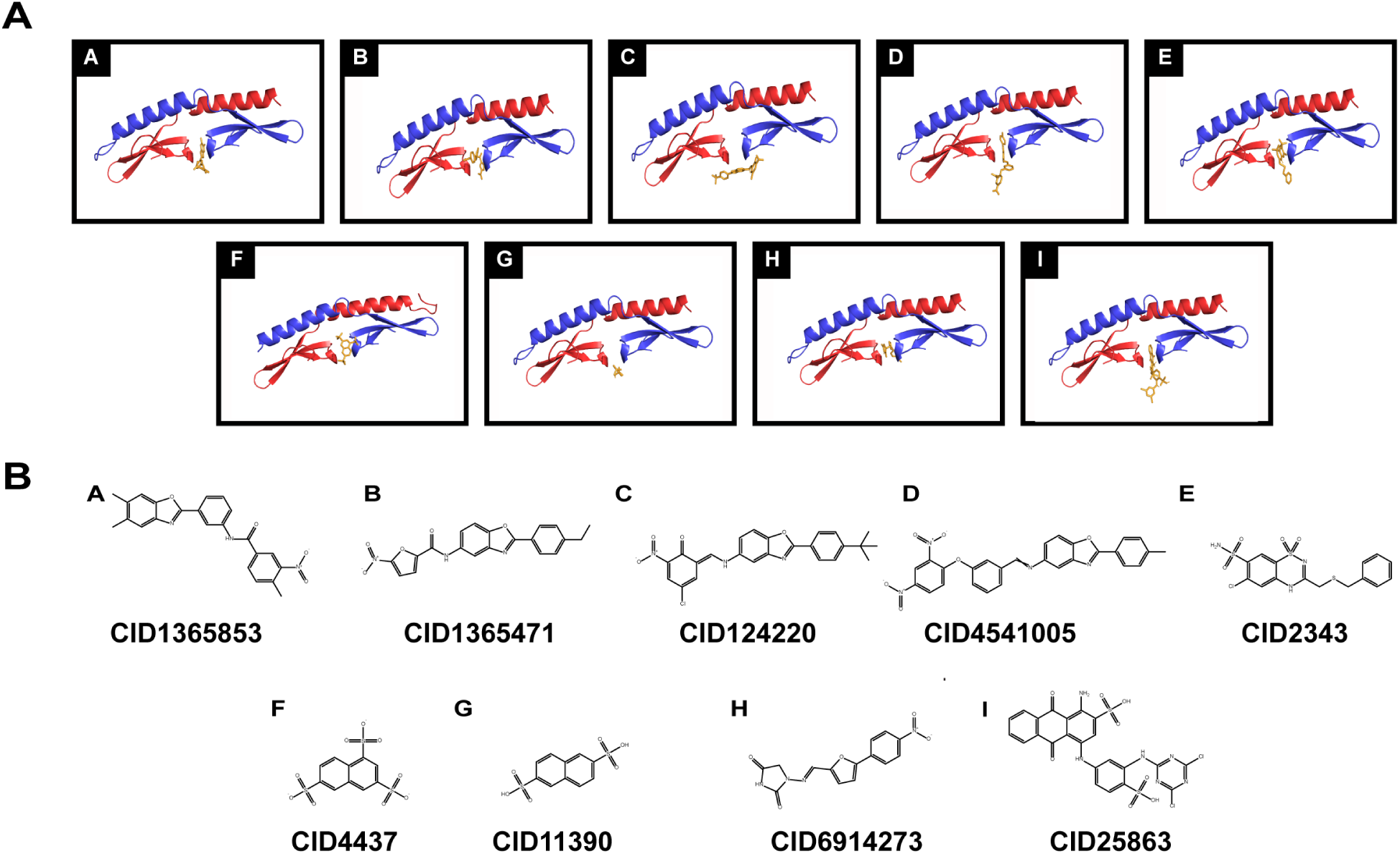
Docking conformations for Compounds A-I. A) The spatial conformation of each of the nine compounds (A-I) that dock within 10 Angstroms of the DNA binding pocket of AP2-EXP with a free energy less than −5kJ/mol is depicted above. B) Chemical structures of each compound (A-I) corresponding to the molecular docking results in panel A.

**Figure S3.**
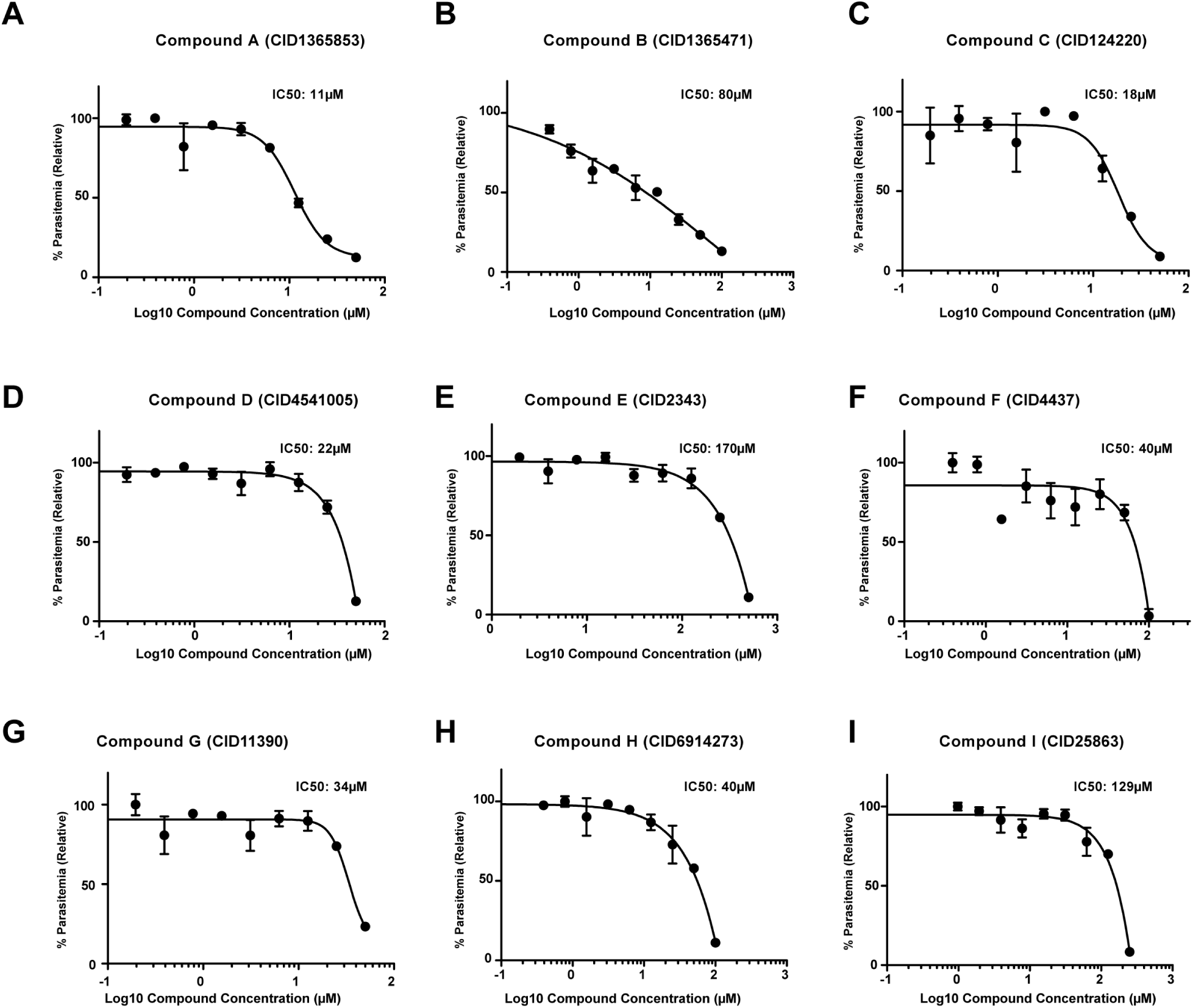
IC50 assays for Compounds A-I. A-I) 48-hour Sybr Green growth inhibition assays were conducted for each of the nine putative ApiAP2 competitor compounds in order to determine IC50 values against asexual *P. falciparum*. All growth assays were performed in triplicate. All compounds kill asexual stage *P. falciparum* parasites in the micromolar (11-170µM) concentration range.

**Figure S4.**
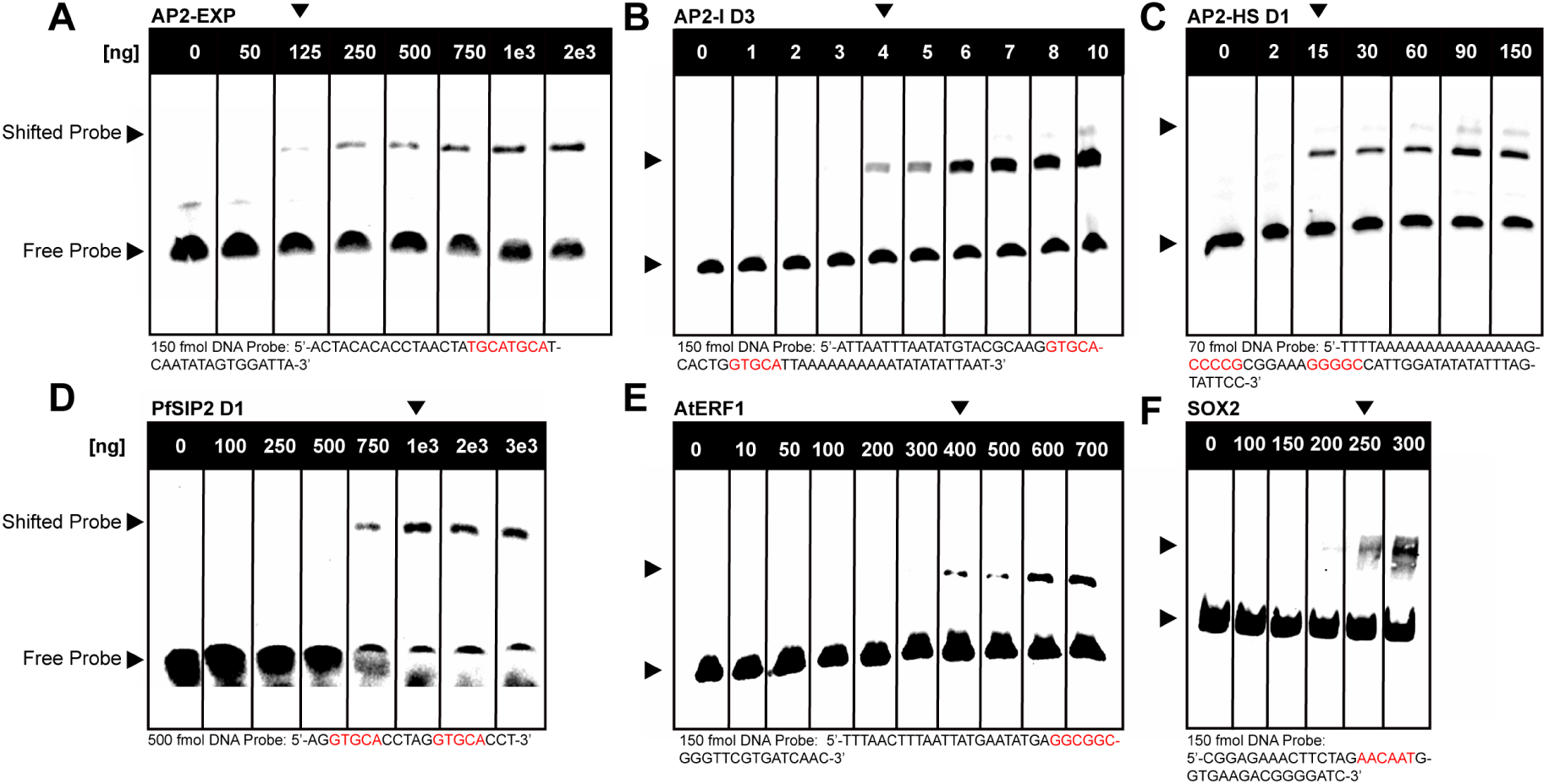
Titration of recombinant DNA binding domains to optimize competition electrophoretic mobility shift (EMSA) assays. A-F) DNA binding domains AP2-EXP, AP2-I D3, AP2-HS D1, PfSIP2 D1, AtERF1, and full length SOX2 were titrated against DNA oligos containing their respective binding motifs (highlighted in red) in an EMSA. Unless otherwise specified, the minimum mass of each recombinant DNA binding domain required to visualize DNA binding (denoted by an arrow) was used in competitive EMSAs with putative ApiAP2 competitor compounds.

**Figure S5.**
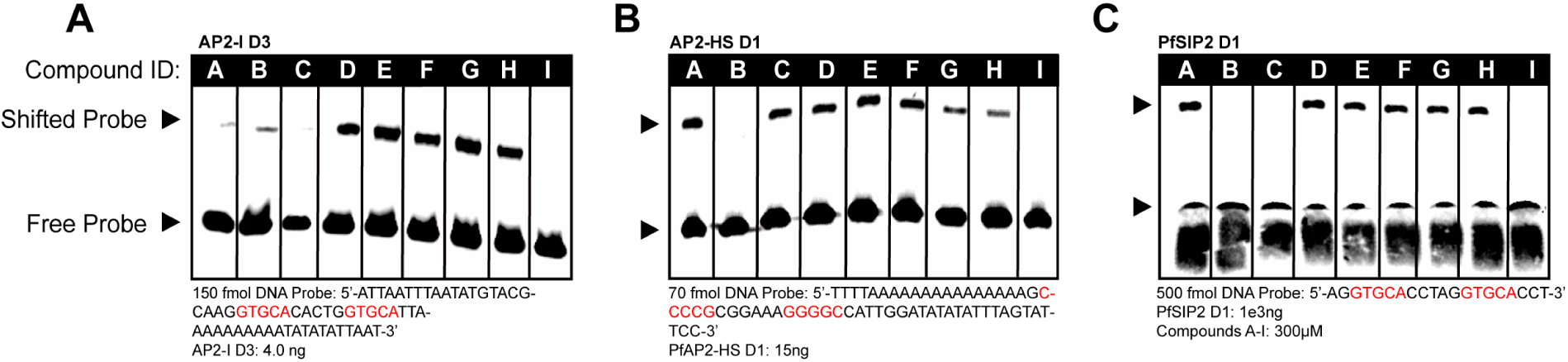
Putative ApiAP2 competitor compounds were tested against three asexual blood stage essential *P. falciparum* AP2 domains in addition to AP2-EXP in a competitive EMSA. AP2-I D3 (A) DNA binding activity is competed by Compounds A, B, C and I. AP2-HS D1 (B) is competed by Compounds B and I, and PfSIP2 D1 (C) is competed by Compounds B, C, and I. Compounds A, B, C and I all compete at least one AP2 domain in addition to AP2-EXP. Cognate DNA motifs for each protein are highlighted in red. 300µM of each compound was used per lane.

**Figure S6.**
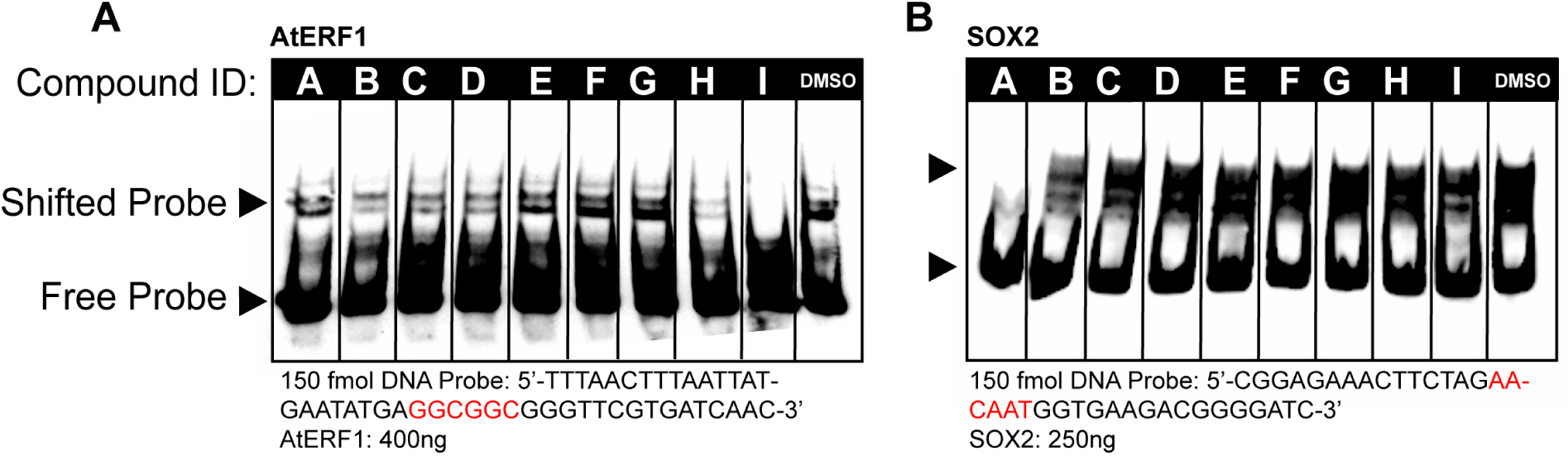
Putative ApiAP2 competitor compounds were tested against off target DNA binding domains in a competitive EMSA. A) The plant encoded *Arabidopsis thaliana* AP2 domain from Ethylene Response Factor 1 (AtERF1) is competed by Compound I. The *Plasmodium* AP2 domain competitors Compound A, B, and C do not compete AtERF1. The cognate AtERF1 DNA motif is highlighted in red. 300µM of each compound was used per lane. B) The human encoded High Mobility Group Box Domain transcription factor SOX2 is competed by Compound A. Due to the lack of homology between SOX2 and AP2 domain proteins, this result indicates that Compound A’s DNA binding competition activity is not unique to the AP2 domain. The cognate SOX2 DNA motif is highlighted in red. 300µM of each compound was used per lane.

**Table S2.**
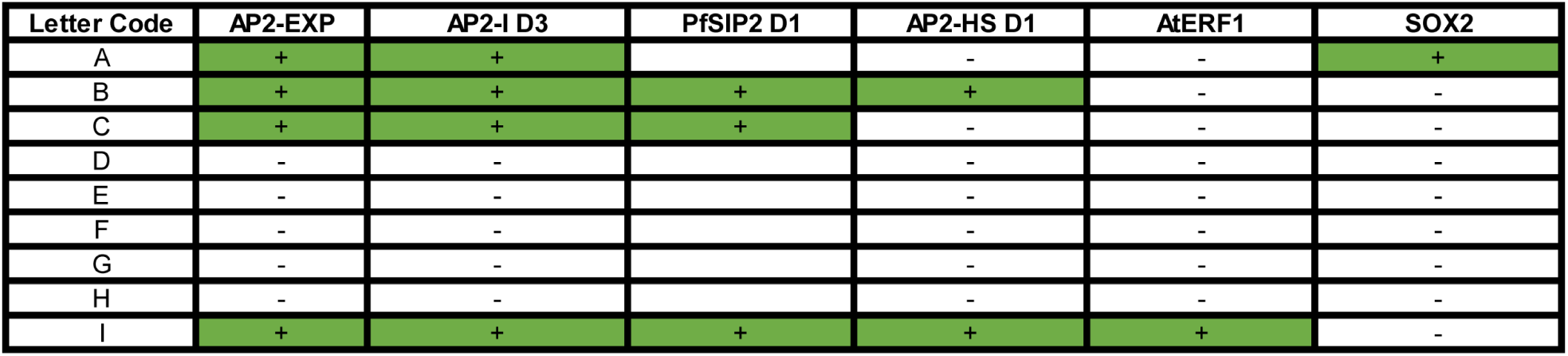
Summary of the Competitive EMSA results for each DNA binding domain tested. Compounds A, B, C, and I all can compete DNA binding by *Plasmodium* AP2 domains. Compounds A and I each compete one off-target protein domain.

**Figure S7.**
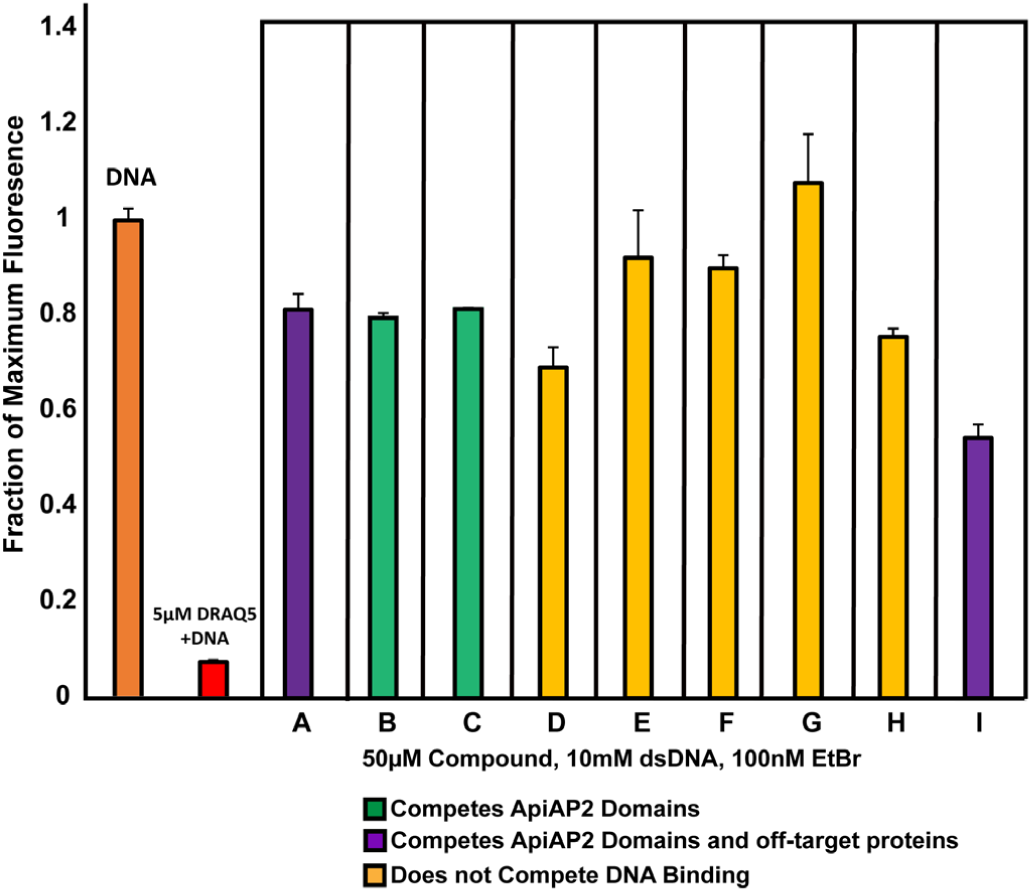
Compounds A-I were tested for DNA intercalation in an ethidium bromide exclusion assay. Each putative ApiAP2 competitor compound was added into a mixture containing double stranded DNA and ethidium bromide. The positive control DNA major groove intercalator DRAQ5 knocks down ethidium bromide fluorescence nearly completely relative to the DNA and ethidium bromide control. The legend indicates the cumulative result for each compound in competitive EMSAs. Each assay was performed in triplicate. Error bars represent standard deviation of the mean.

**Figure S8.**
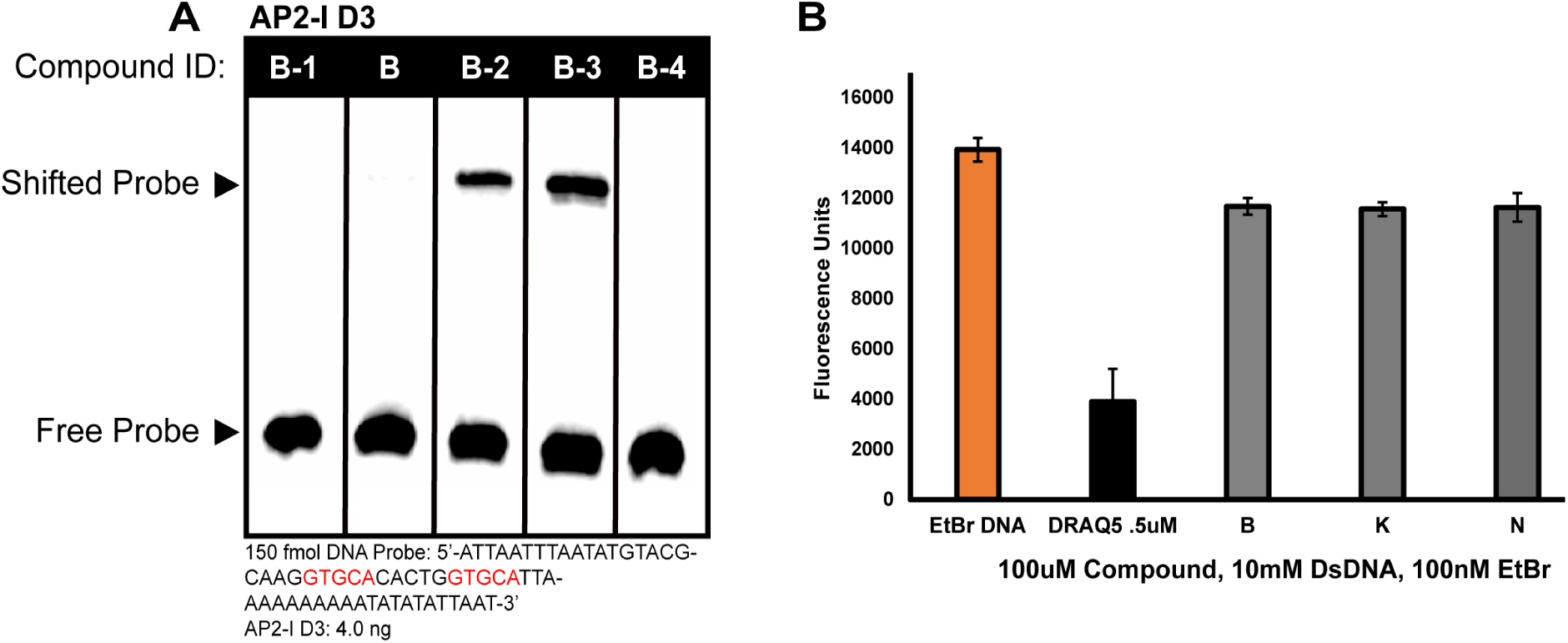
Compound B analogues were tested against AP2-I D3 in a competitive EMSA and checked for DNA intercalation ability. A) Compounds B, B-1, B-2, B-3, and B-4 were added to an EMSA with AP2-I D3 to check whether their DNA binding competition is consistent with AP2-EXP. The cognate AP2-I D3 DNA motif is highlighted in red. 300µM of each compound was used per lane. B) Compounds B, B-1, B-2, B-3, and B-4 were tested for DNA major groove intercalation in an ethidium bromide exclusion assay. DRAQ5 was used as a positive control for intercalation. Each assay was performed in triplicate. Error bars represent standard deviation of the mean.

**Movies S1-S5 Molecular dynamics simulations for Compound B and Compound B analogues B-1, B-2, B-3, and B-4 interaction with AP2-EXP**

Compound B (Movie S1), Compound B-1 (Movie S2), Compound B-2 (Movie S3), Compound B-3 (Movie S4), and Compound B-4 (Movie S5) interactions with AP2-EXP were each simulated using molecular dynamics. Each molecule was initially started in the location predicted for Compound B by molecular docking.

**Figure S9.**
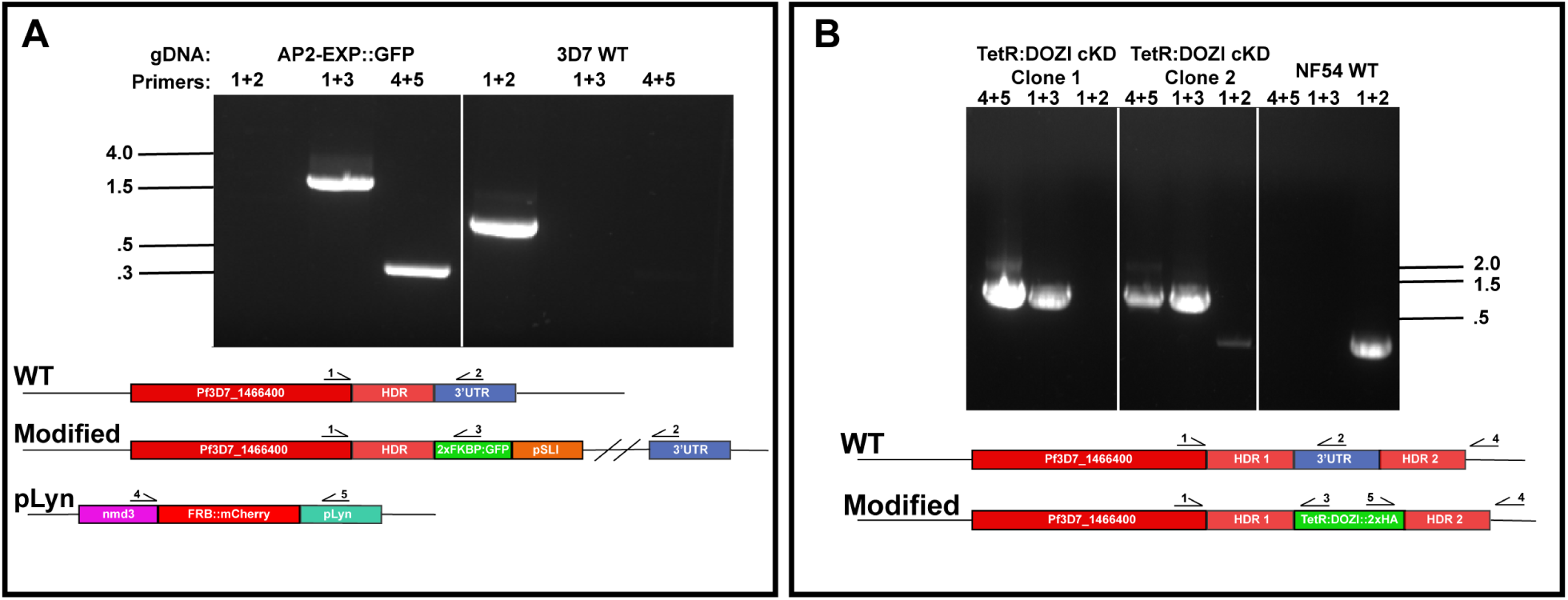
Creation of endogenously tagged parasite lines AP2-EXP::GFP and AP2-EXP::HA. A) The Selection Linked Integration system was used to add 2xFKBP inducible mislocalization protein and GFP to AP2-EXP by single homologous recombination. Successful integration was confirmed by genotyping PCR. Genomic DNA from the wild type Pf3D7 parental line was used as a control. In order to test the efficacy of the knock sideways system, the pLyn mislocalizer plasmid was added to AP2-EXP::GFP and confirmed by PCR. DNA kb are indicated by the marks to the left of the gel. B) The PSN054 TetR:DOZI plasmid was used to add the TetR:DOZI mRNA repression module and endogenous 2xHA tag to AP2-EXP by double homologous recombination. Correct integration was confirmed by genotyping PCR. The parental NF54 parasite line was used as the unedited control. Clonal populations one and two are indicated as C1 and C2, respectively. Clone one had correct integration and complete absence of the wild type *ap2-exp* DNA locus and was used for further experiments. DNA kb are indicated by the marks to the right of the gel.

**Figure S10.**
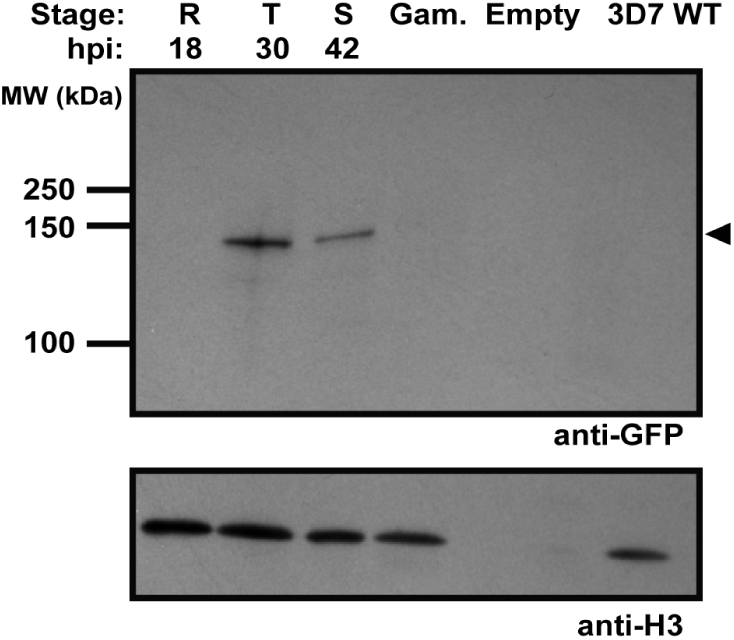
AP2-EXP protein expression in the AP2-EXP::GFP endogenously tagged parasite line (related to figure 3A) AP2-EXP expression was tracked throughout the IDC by harvesting protein from highly synchronous asexual blood stage parasites followed by a western blot against the GFP tag. Histone H3 was used as a loading control.

**Figure S11.**
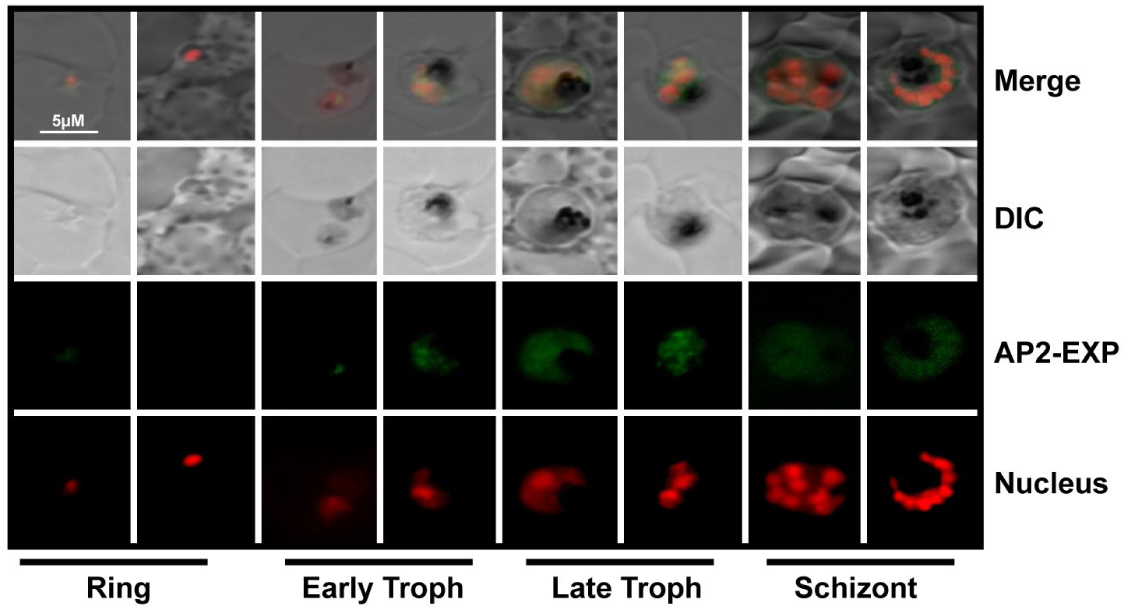
Fluorescent microscopy tracking of AP2-EXP::GFP expression. AP2-EXP::GFP expression was monitored in a highly synchronous parasite population by fluorescent microscopy across the IDC. DRAQ5 was used as a nuclear stain for parasites.

**Figure S12.**
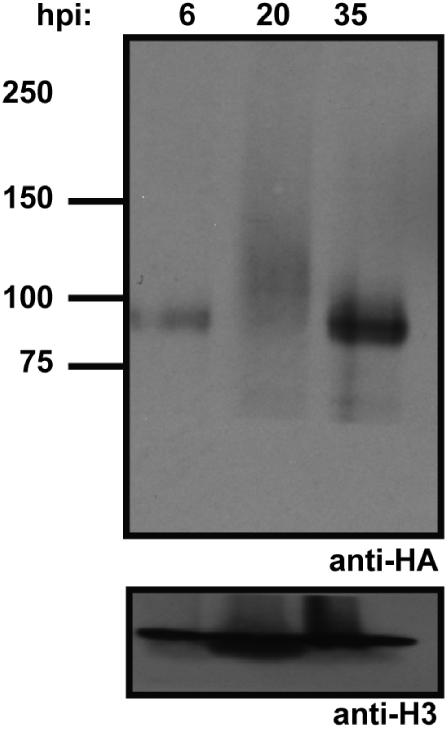
AP2-EXP protein expression in the AP2-EXP::HA endogenously tagged parasite line. AP2-EXP expression was tracked throughout the IDC by harvesting protein from highly synchronous asexual blood stage parasites followed by a western blot against the 2xHA tag. Histone H3 was used as a loading control.

**Figure S13.**
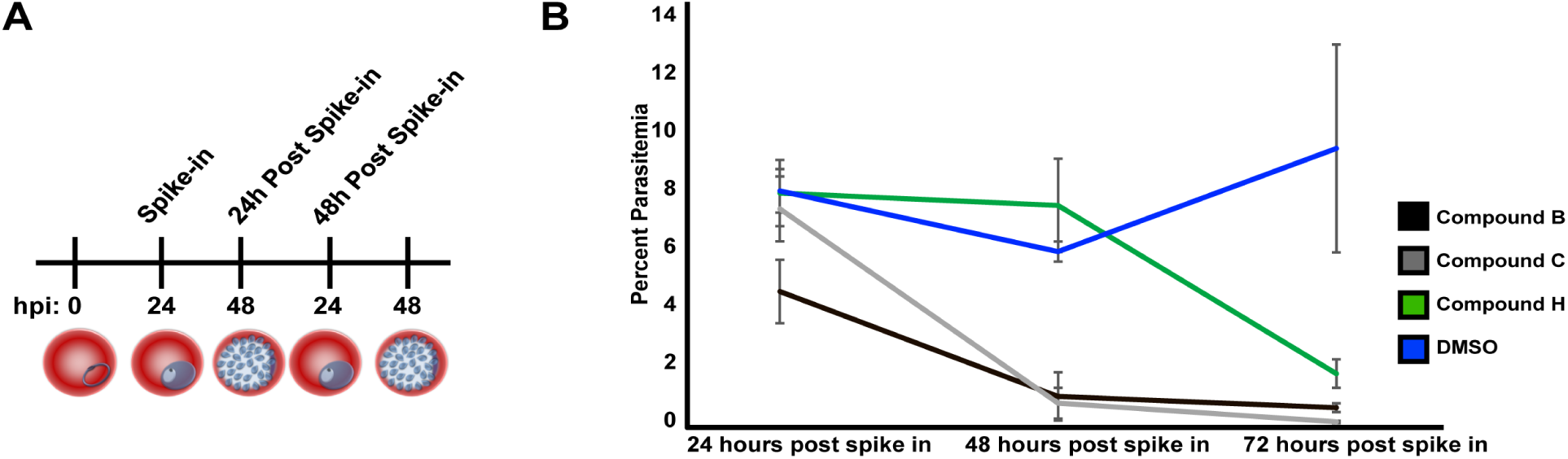
Parasite counts for 48-hour Compounds B, C, and H phenotyping time course, related to figure 3B. A) Schematic for AP2 domain competitor compound phenotyping time course. B) 40µM of ApiAP2 competitor Compounds B or C, or non ApiAP2 competitor Compound H, was spiked into highly synchronous wild type Pf3D7 parasites at 24 hours post invasion. DMSO vehicle was used as a reference for normal growth. All growth assays were performed in triplicate. Error bars represent standard deviation of the mean.

**Figure S14.**
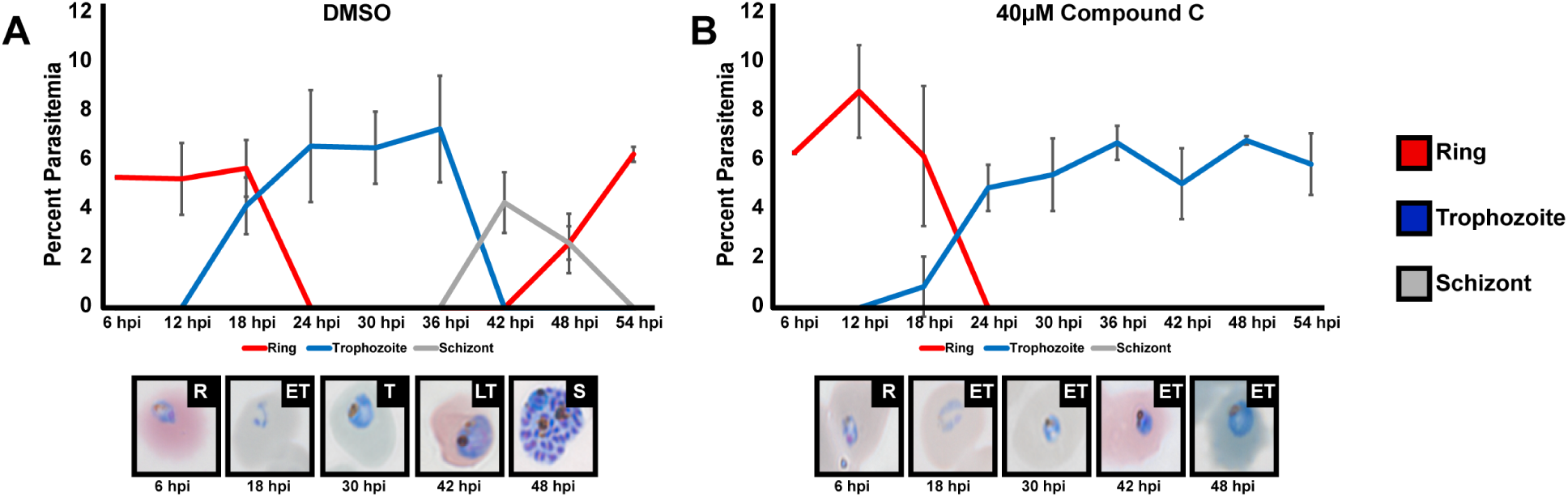
Compound C causes parasites to stop progressing in the early trophozoite stage at 30 hpi. Highly synchronous wild type Pf3D7 parasites were spiked with DMSO vehicle control (A) or 40µM Compound C (B) and monitored at 6-hour intervals throughout the IDC. Each growth assay was performed in triplicate. Error bars represent standard deviation of the mean. Giemsa stained parasite images are representative of each population at the specified time point. R, ET, T, LT, and S indicate ring, early trophozoite, late trophozoite, or schizont morphologies, respectively.

**Figure S15.**
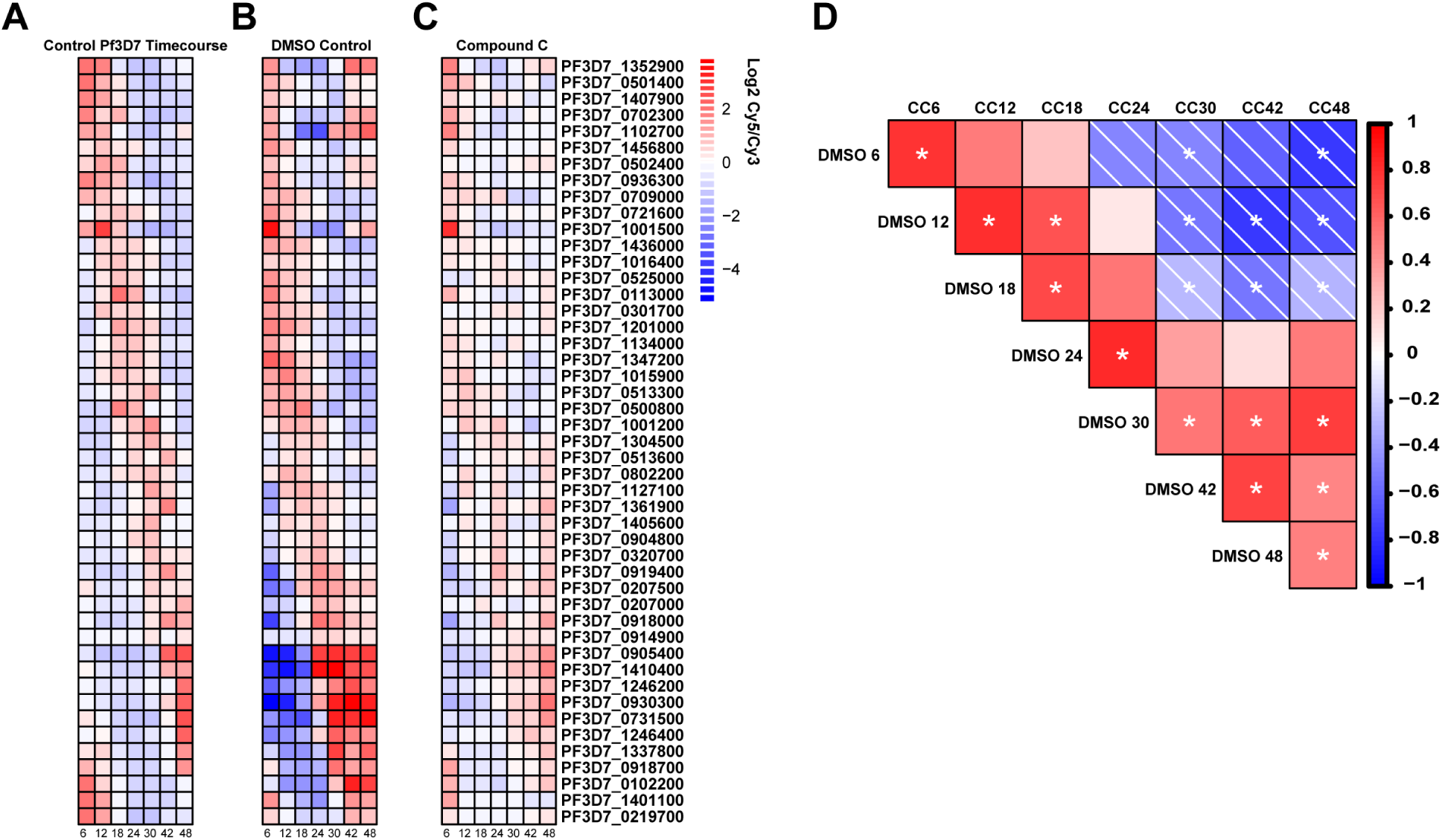
Quality control of DNA microarray data for DMSO vehicle control and Compound C parasites, related to figure 4. A) DNA microarray data from(34) for a set of highly periodic control genes expressed in the IDC. B-C) The same set of highly periodic control genes as in panel A were plotted for the DMSO control and 12µM Compound C spiked parasites in order to compare parasite staging between the two experiments. D) Correlogram depicting the Spearman Correlation value between control gene expression for DMSO (Panel B) and Compound C (Panel C) samples. A * indicates p value < .05.

**Figure S16.**
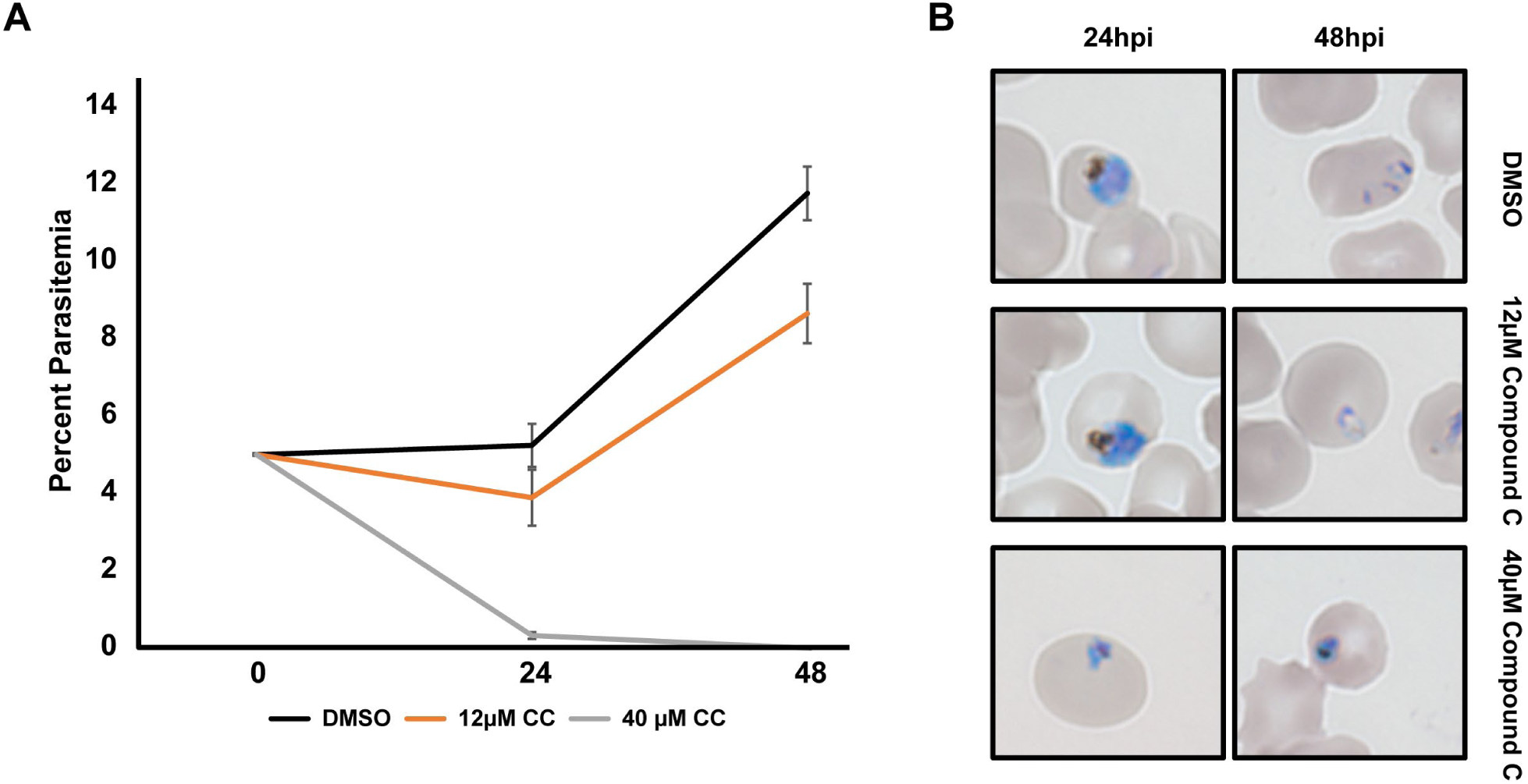
A 48-hour time course to determine the phenotype for 12µM Compound C, related to Figure 4A. A) Highly synchronous asexual blood stage Pf3D7 parasites were spiked with DMSO vehicle control, 12µM Compound C, or 40µM Compound C. 40µM Compound C was used as a positive control for complete parasite death at 48 hpi. Each growth assay was performed in triplicate. Error bars represent standard deviation of the mean. B) Representative images of each parasite population (DMSO, 12µM Compound C, 40µM Compound C) at 24 and 48 hpi.

**Figure S17.**
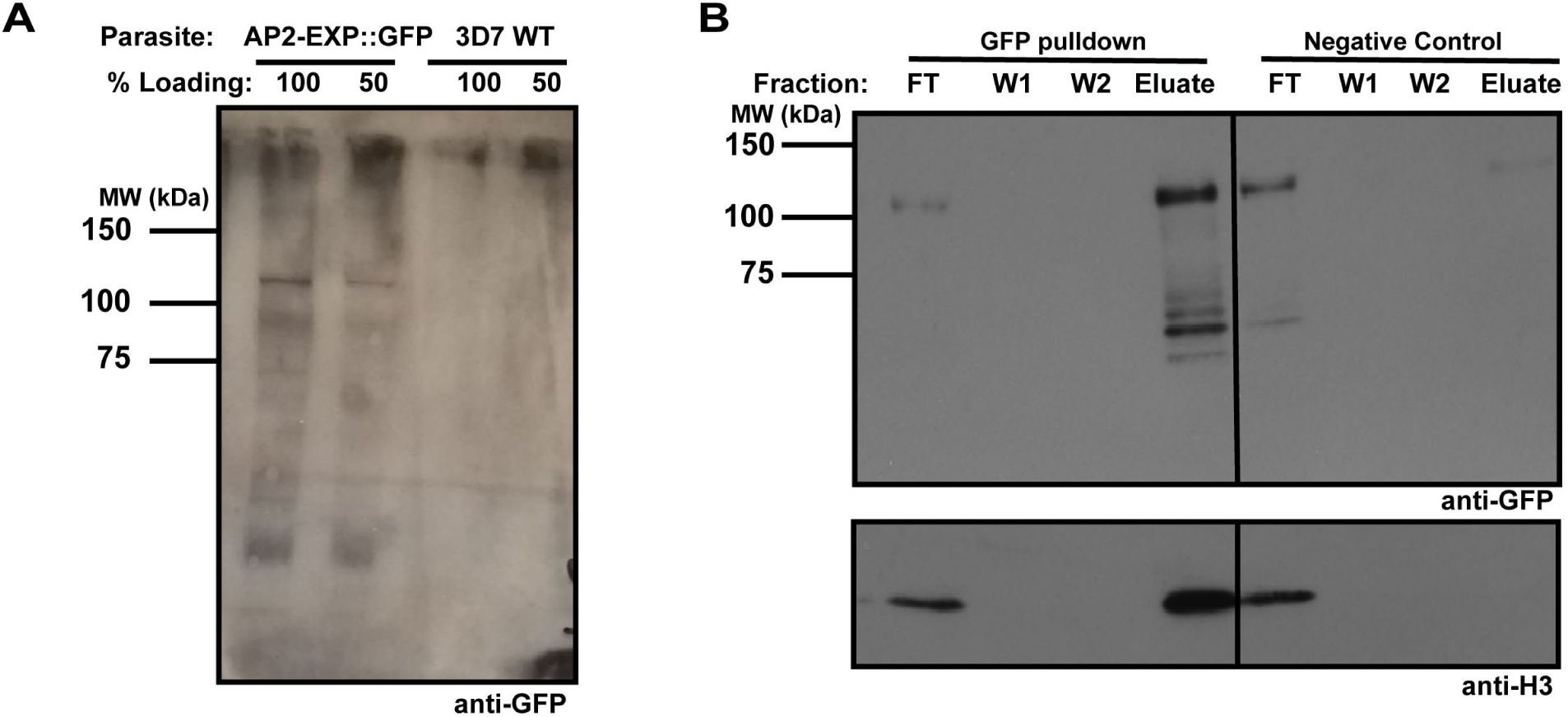
ChIP-seq protein quality control, related to figure 4B. A) Crosslinked nuclear material was blotted after sonication to ensure recovery of the full-length AP2-EXP protein during chromatin immunoprecipitation. Full length AP2-EXP is recovered, indicating that the protocol is suitable to analyze AP2-EXP DNA binding *in vivo*. Crosslinked nuclear material from the wildtype Pf3D7 parental parasite line was used as a negative control. B) Anti-GFP beads were used to pull down GFP tagged AP2-EXP from AP2-EXP::GFP. Flowthrough (FT), Wash (W1 and W2) and Eluate fractions were saved and analyzed by western blot. The presence of AP2-EXP and Histone H3 in the Eluate lane indicates that AP2-EXP interacts with chromatin in the nucleus. The non-immune negative control beads do not enrich AP2-EXP or Histone H3 in the eluate.

**Figure S18.**
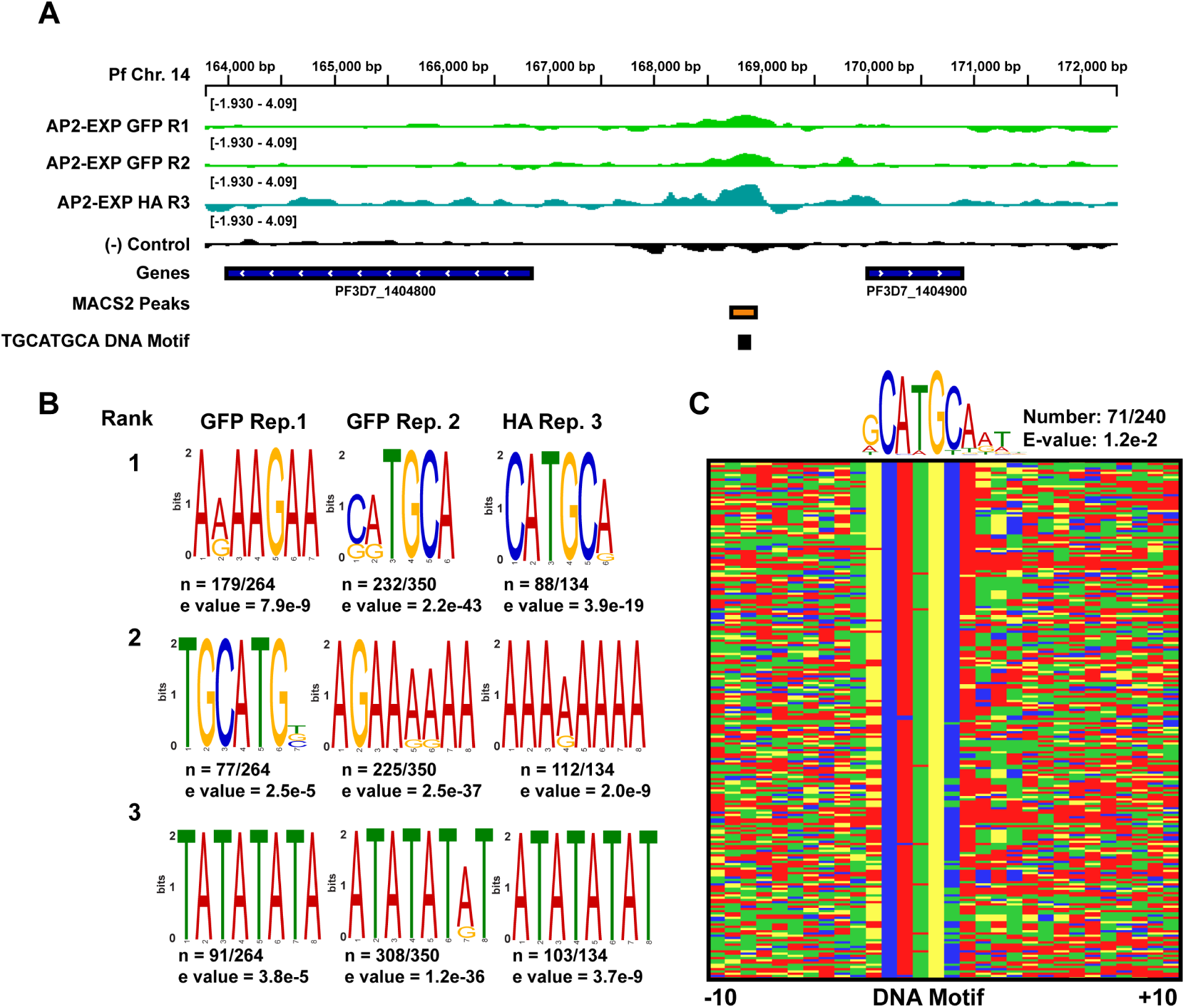
ChIP-seq extended data. A) Log2 immunoprecipitate/input ChIP-seq data from each replicate (2x AP2-EXP::GFP and 1xAP2-EXP::HA) of AP2-EXP ChIP-seq visualized by IGV at a representative DNA locus. The location of a conserved MACS2 called peak of occupancy and the TGCATGCA DNA motif is indicated by the bottom tracks. The (-) control lane is the coverage resulting from a no-epitope control ChIP-seq. B) The top three ranked DNA motifs present within peaks of occupancy for each ChIP-seq replicate as determined by DREME(24). The core DNA motif CATGCA is overrepresented within each individual replicate. C) The top overrepresented DNA motif within AP2-EXP peaks of occupancy conserved in 2/3 replicates of ChIP-seq as determined by DREME(24) plotted at the primary DNA sequence level. DNA sequences were sorted from highest to lowest degree of motif conservation using FIMO.(25)

**Figure S19.**
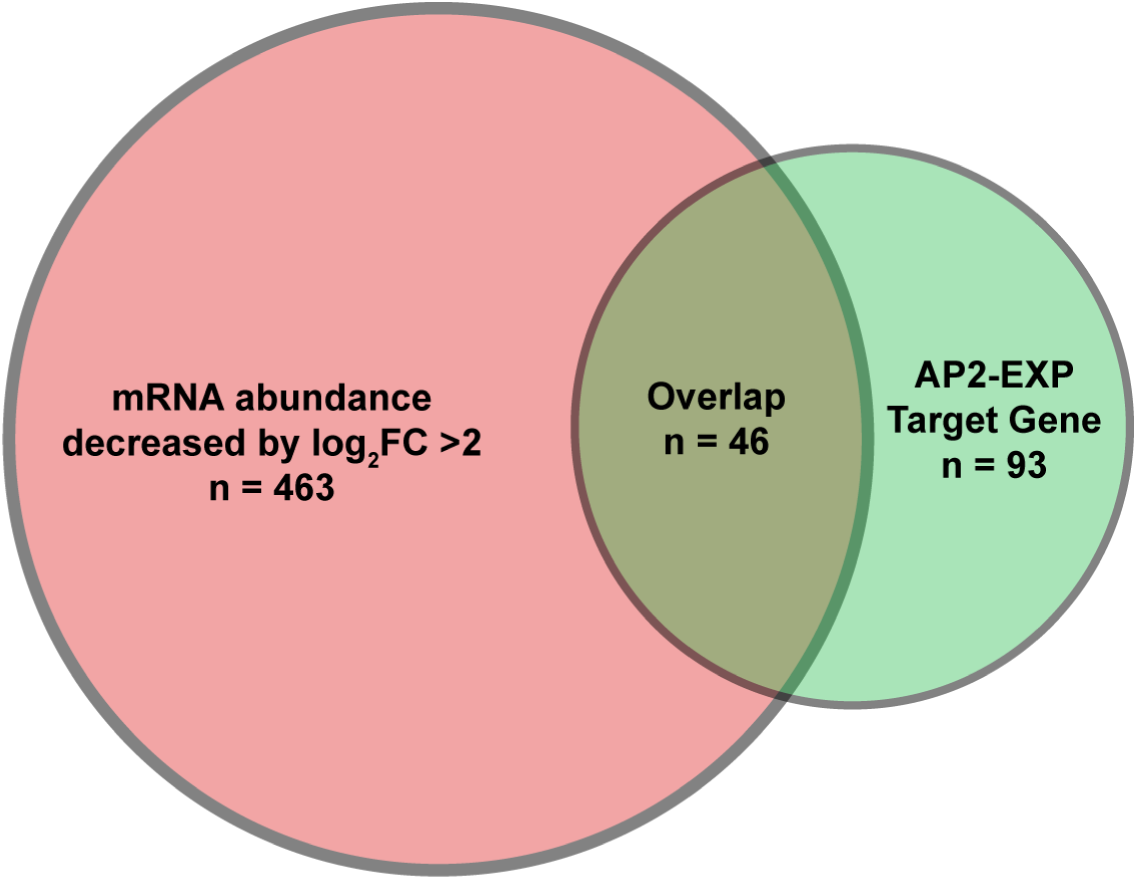
Comparison of AP2-EXP target genes with Compound C induced changes in transcript abundance. The total overlap between AP2-EXP gene targets detected in the Compound C RNA time course and global decrease in transcript abundance at 24-30 hpi by log_2_ fold change >2.

**Figure S20.**
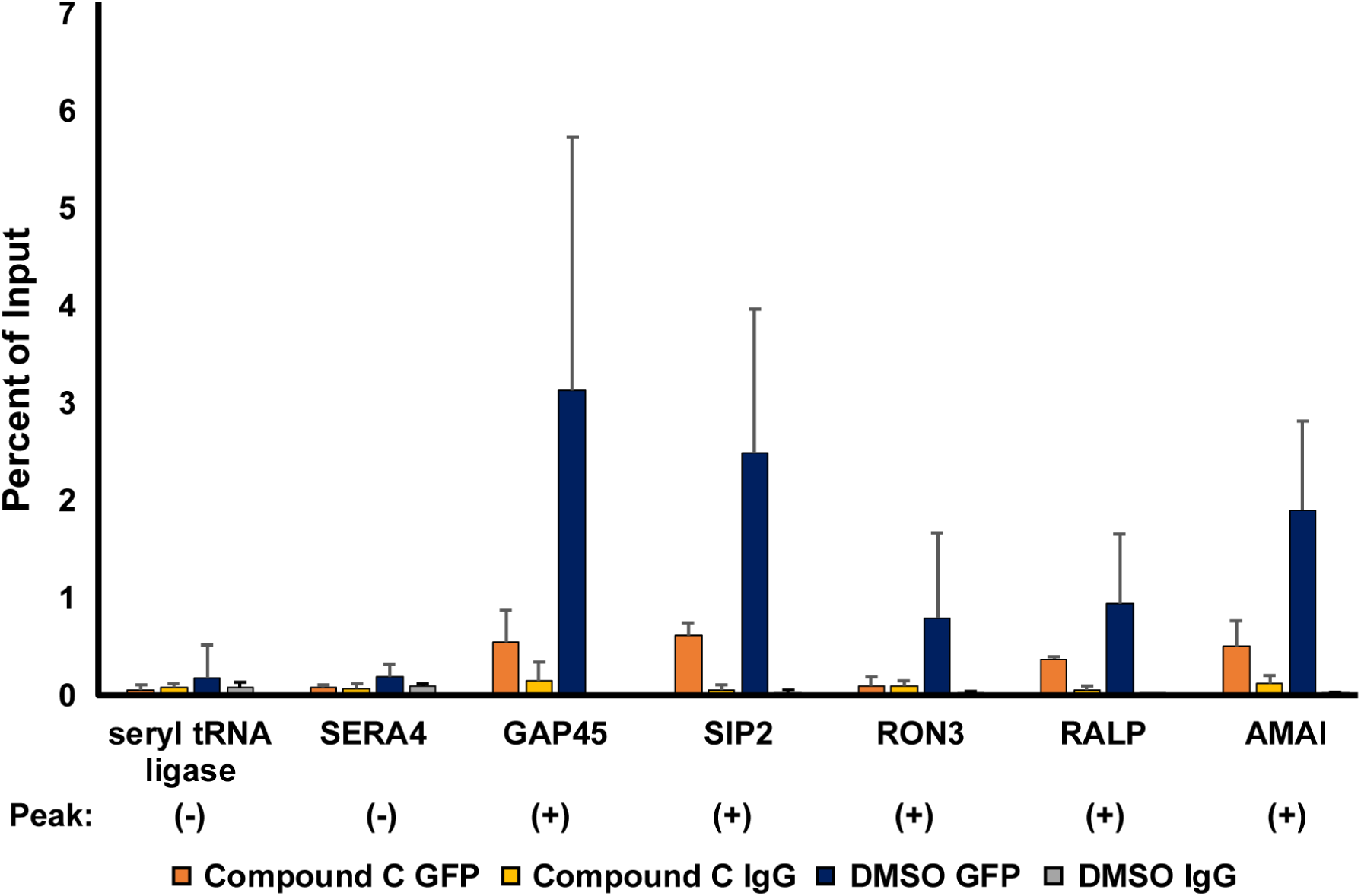
ChIP-Quantitative PCR to assess Compound C impact on AP2-EXP genomic occupancy. AP2-EXP::GFP parasites were spiked with 40µM Compound C or DMSO vehicle control at 30 hpi for two hours. ChIP samples were collected for each population using either anti-GFP or negative control IgG antibodies. The percent of input was determined by RT-qPCR. Each experiment was performed in triplicate. The presence or absence of an AP2-EXP peak of occupancy at each DNA locus based on ChIP-seq is indicated by a (+) or (-), respectively. Each assay was done in triplicate with the exception of AMAI, where n =2.

**Figure S21.**
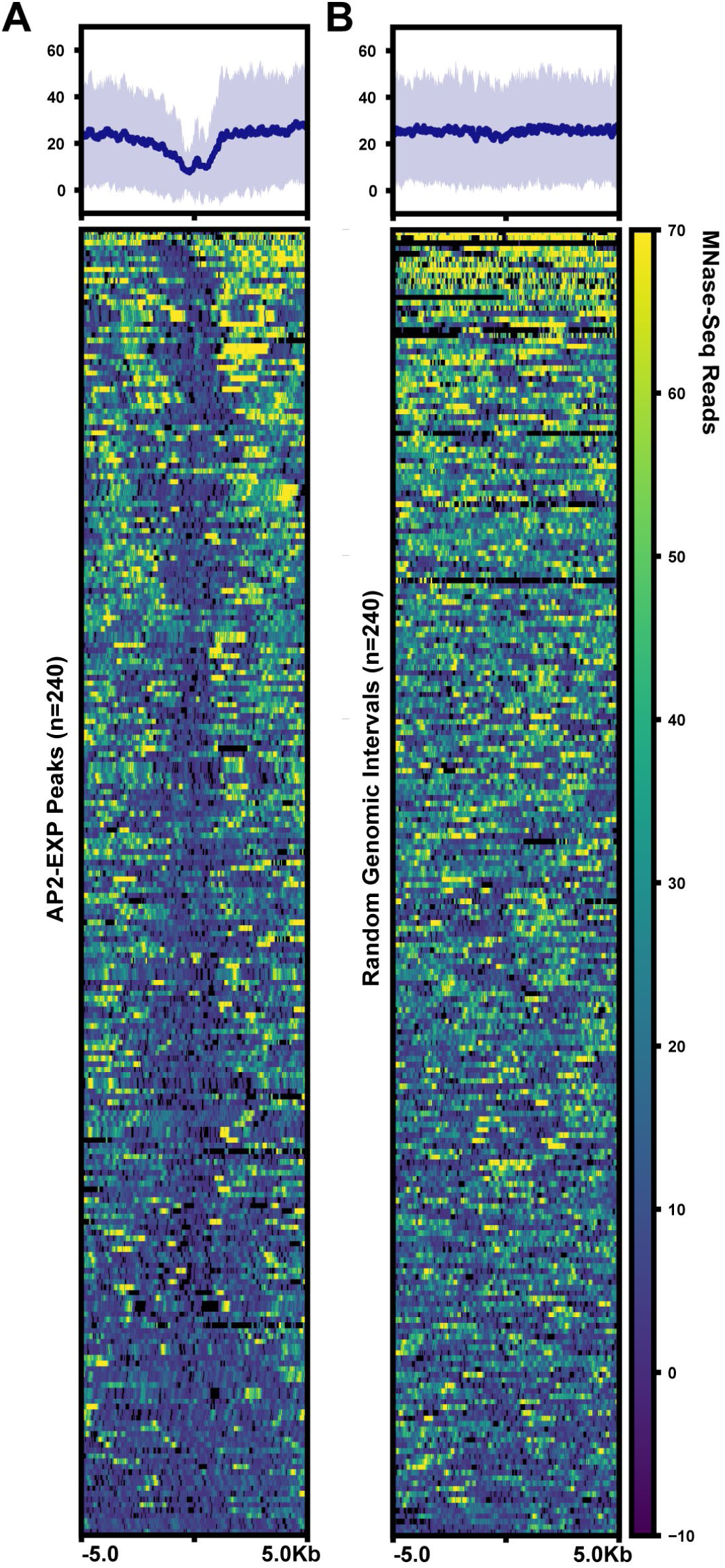
Nucleosome occupancy is depleted at AP2-EXP DNA binding sites. Mnase-seq data(30) was plotted against DNA binding sites conserved in 2/3 replicates of AP2-EXP ChIP-seq (A) or random genomic intervals of equal length from the same chromosome as the original peak (B).

**Figure S22.**
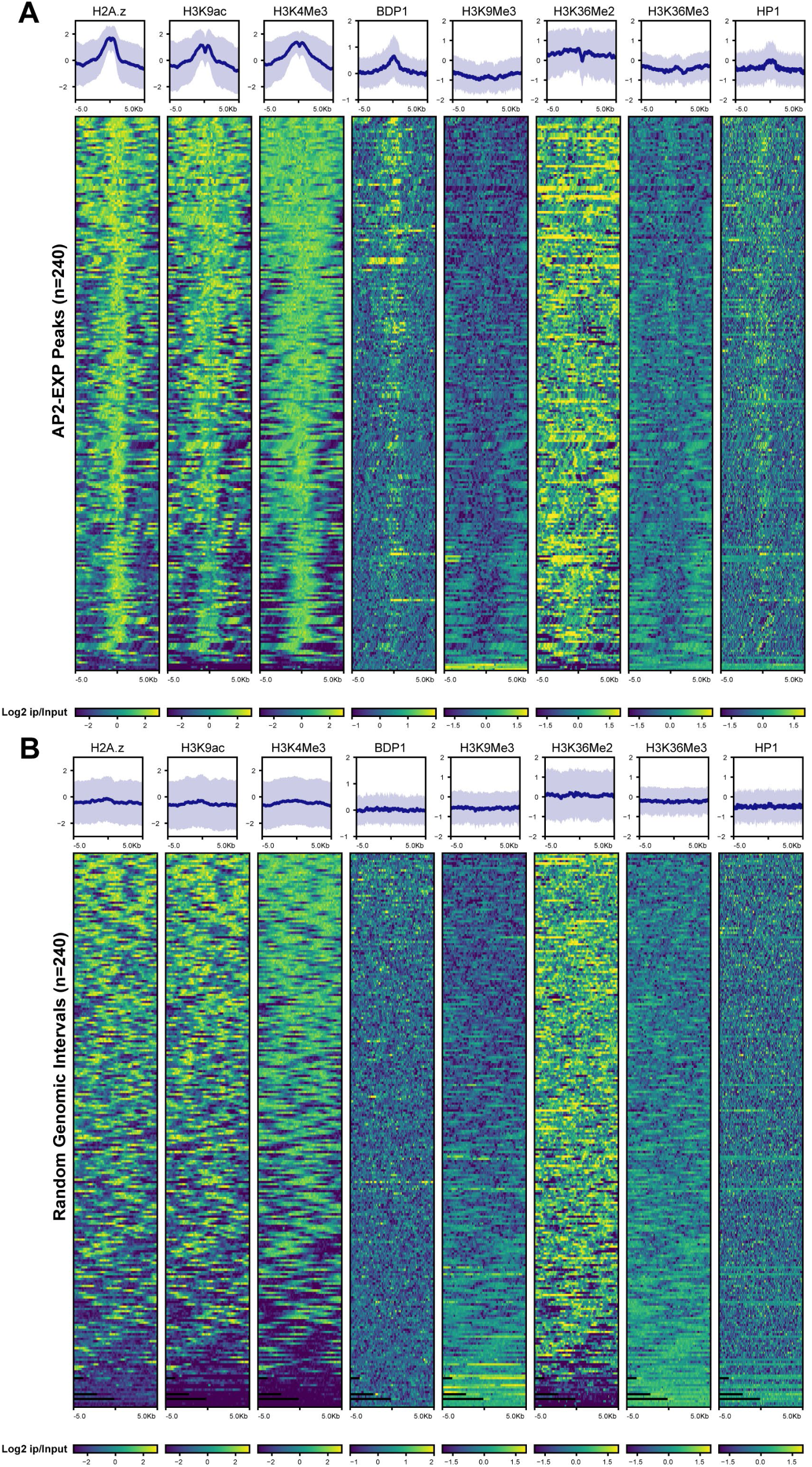
Histone post translational modifications and chromatin reader occupancy at AP2-EXP peaks. The occupancy of histone variant H2A.z, histone modifications H3K9ac, H3K4me3(27), H3K9me3, H3K36me2/3(26), and chromatin readers BDP1(29) and HP1(31) were plotted against AP2-EXP peaks of occupancy conserved in 2/3 ChIP-seq replicates (A) or random genomic intervals of the same length (B), taken from the same chromosome on which the AP2-EXP peak originally occurred.

**Figure S23.**
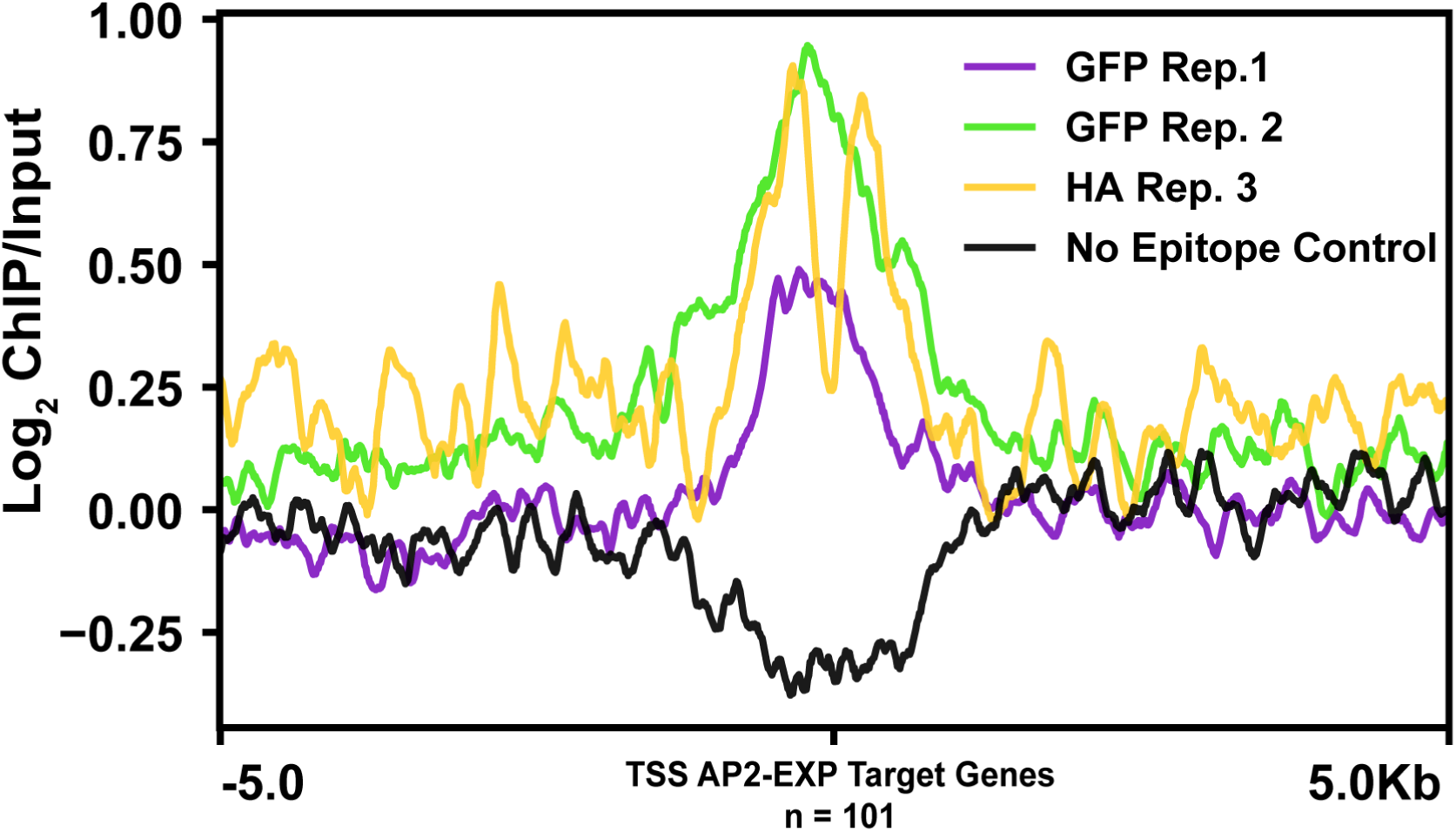
AP2-EXP DNA occupancy with respect to the Transcription Start Site (TSS) of target genes. Log_2_ immunoprecipitate (ChIP)/Input ChIP-seq coverage for each replicate of AP2-EXP ChIP-seq and the no epitope control was plotted against the TSS(35) of each target gene conserved in 2/3 ChIP-seq replicates.

**Figure S24.**
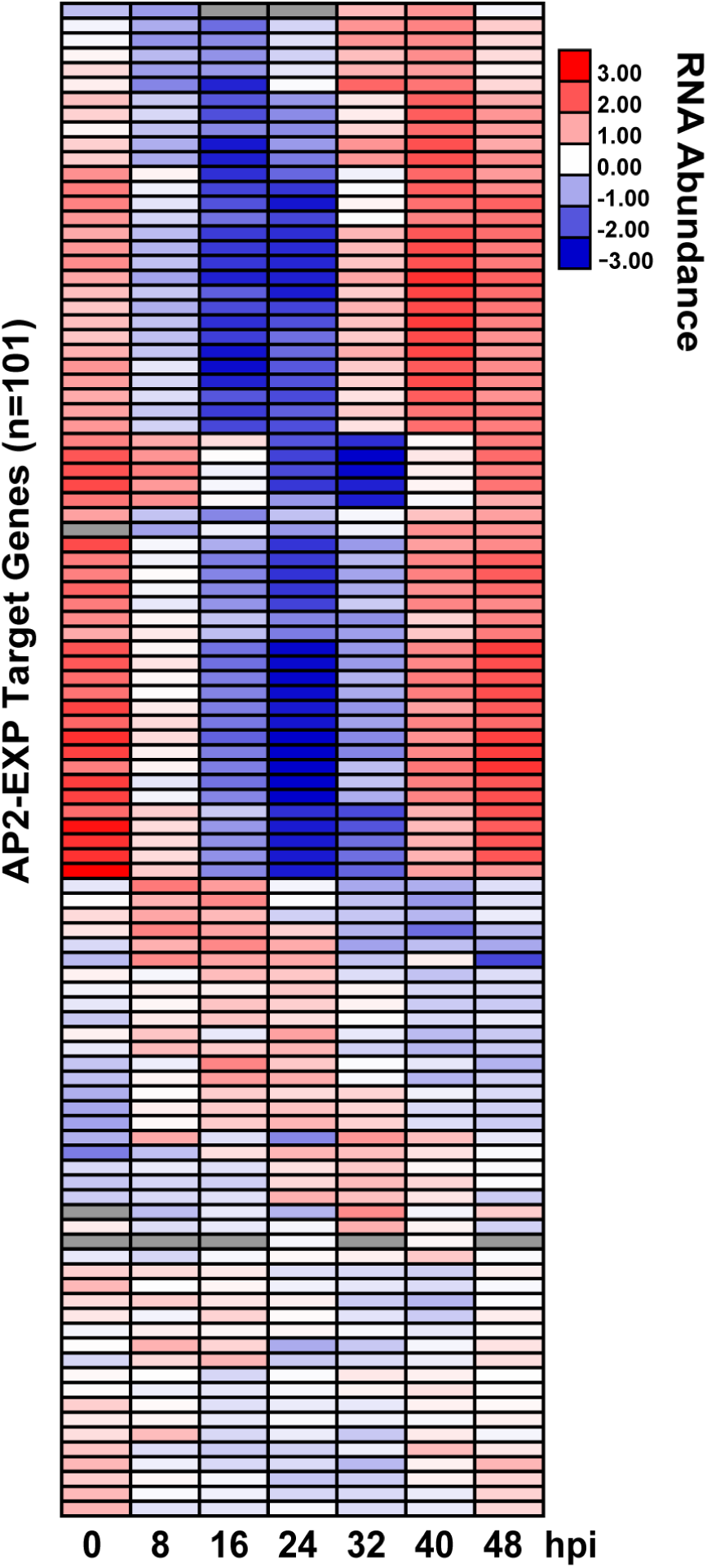
Normal Transcript Abundance of AP2-EXP Target Genes. AP2-EXP target genes were determined by ChIP-seq and their transcript abundance data during the 48-hour IDC was plotted using data from Chappell *et al*(35).

**Figure S25.**
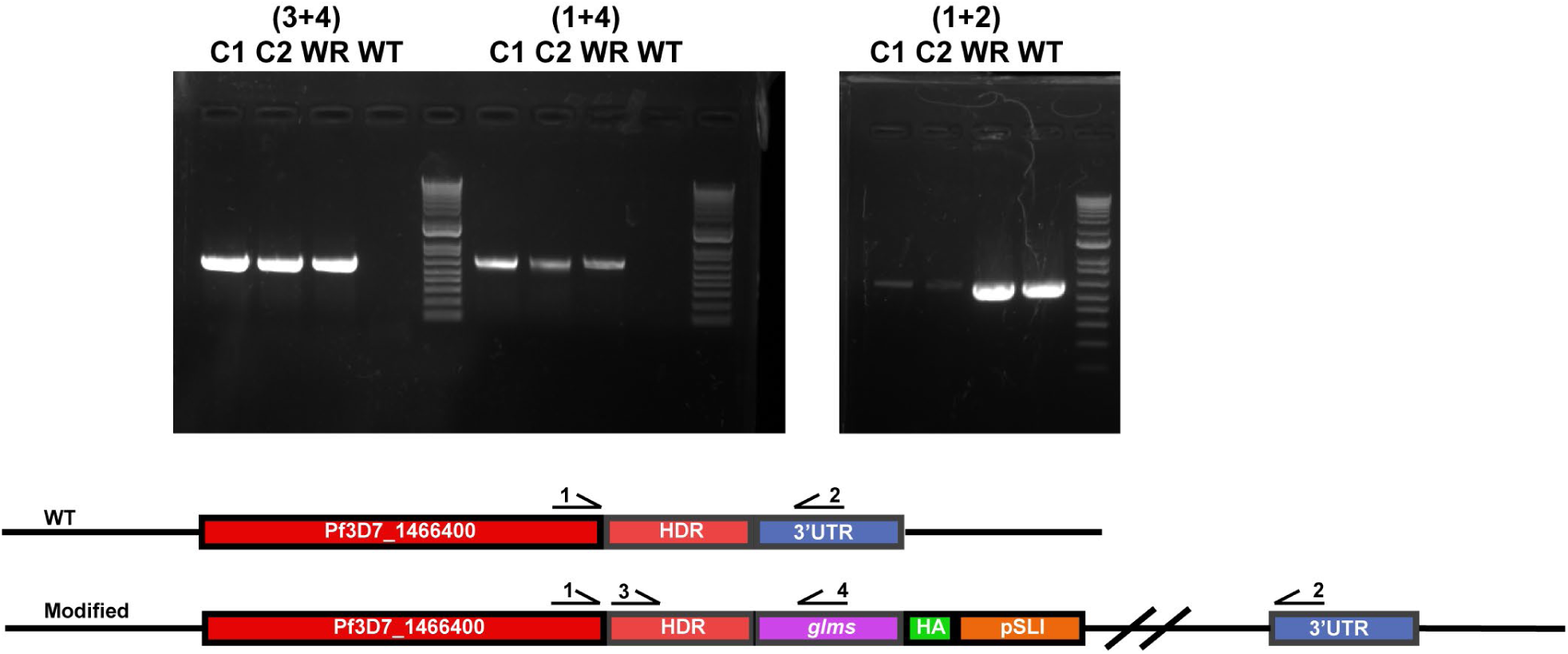
Creation of a *glms* ribozyme based knockdown line for AP2-EXP. Wildtype Pf3D7 *Plasmodium falciparum* parasites were transfected with pSLI:AP2-EXP*::glms*::HA to endogenously tag the AP2-EXP DNA locus with an inducible *glms* ribozyme and HA epitope tag. Successful integration to create AP2-EXP::*glms*::HA by single crossover homologous recombination was confirmed by genotyping PCR. C1 and C2 represent clonal populations selected for integration. WR represents a parasite population selected for the plasmid but not for integration. WT represents Pf3D7 wild type control parasite gDNA.

**Figure S26.**
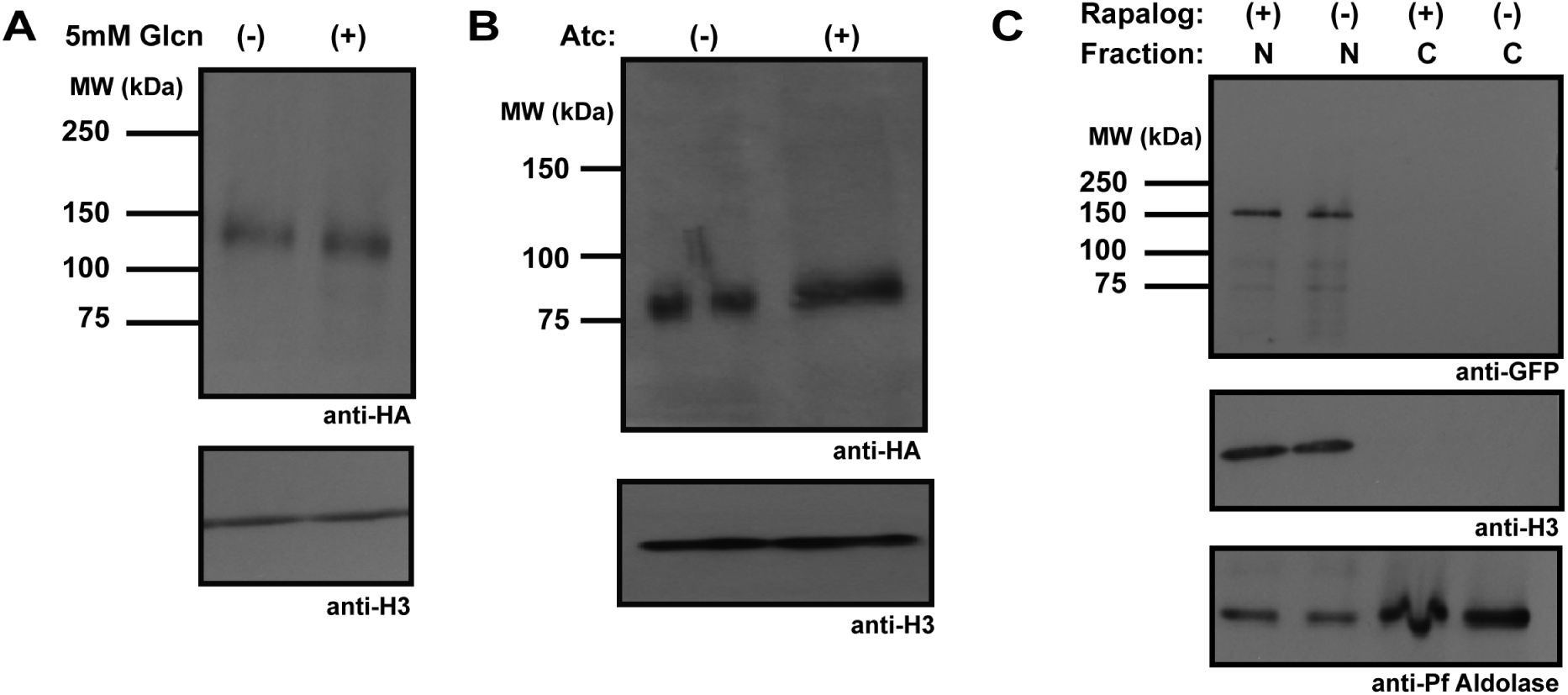
Western blot phenotyping of attempts to genetically knockdown AP2-EXP. A) To assess genetic knockdown of AP2-EXP by *glms* ribozyme tag in the parasite line AP2-EXP::*glms*::HA, highly synchronous parasites were spiked with 5mM glucosamine or vehicle control for 72 hours and AP2-EXP quantity was determined by anti-HA western blot. Histone H3 was used as a loading control. Glucosamine treatment did not impact the amount of AP2-EXP protein present. B) Genetic knockdown of AP2-EXP by the TetR:DOZI mRNA repression module was assessed in the parasite line AP2-EXP::HA by washing anhydrotetracycline (aTc) out of the media for 72 hours. AP2-EXP quantity was determined by anti-HA western blot, and Histone H3 was used as a loading control. Removal of aTc from the media did not impact the amount of AP2-EXP protein present. C) Genetic knockdown of AP2-EXP via protein mislocalization was assessed for the parasite line AP2-EXP::GFP. 250nM rapalog was added to the media for 48 hours and AP2-EXP protein localization was assessed by ant-GFP western blot. Histone H3 and Aldolase were used as nuclear and cytosolic markers, respectively. The addition of rapalog did not cause any detectable mislocalization of AP2-EXP from the nucleus to the cytosol. N indicates the nuclear protein fraction, and C indicates the cytosolic fraction.

**Figure S27.**
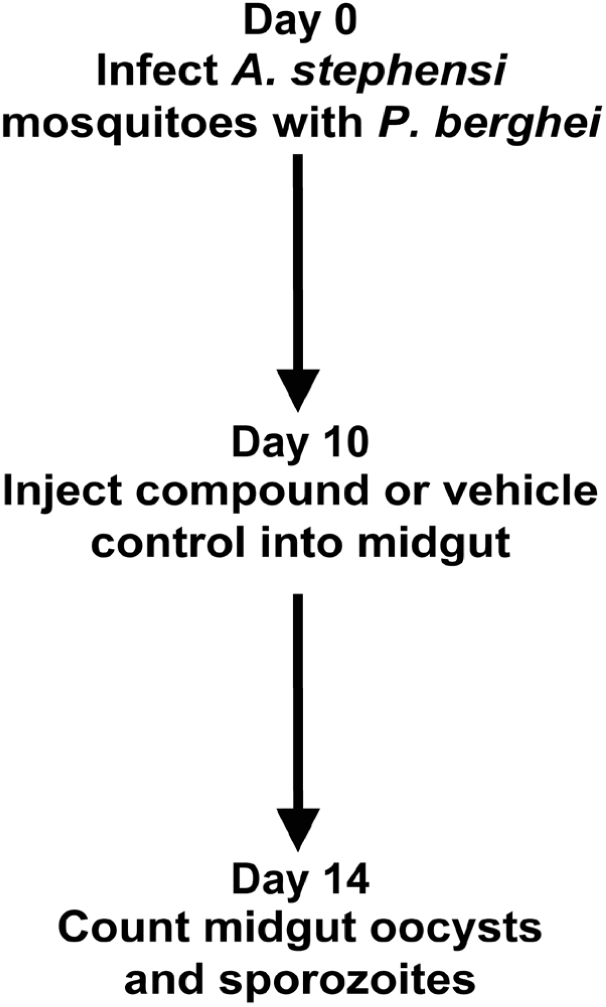
Mosquito stage *P. berghei* inhibition assay schematic. *A. stephensi* mosquitoes were infected with *Plasmodium berghei* parasites. On day 10 post infection, mosquito midguts were injected with the measured IC50 of Compounds B, C, F, or DMSO vehicle control. On day 14 post infection mosquitoes were dissected to count oocysts and midgut sporozoites. For Compound C and DMSO vehicle control, each experiment was performed in duplicate. Compounds B and F phenotyping were performed as a single experiment.

